# Sensei: How many samples to tell evolution in single-cell studies?

**DOI:** 10.1101/2020.05.31.126565

**Authors:** Shaoheng Liang, Jason Willis, Jinzhuang Dou, Vakul Mohanty, Yuefan Huang, Eduardo Vilar, Ken Chen

## Abstract

Cellular heterogeneity underlies cancer evolution and metastasis. Advances in single-cell technologies such as single-cell RNA sequencing and mass cytometry have enabled interrogation of cell type-specific expression profiles and abundance across heterogeneous cancer samples obtained from clinical trials and preclinical studies. However, challenges remain in determining sample sizes needed for ascertaining changes in cell type abundances in a controlled study. To address this statistical challenge, we have developed a new approach, named Sensei, to determine the number of samples and the number of cells that are required to ascertain such changes between two groups of samples in single-cell studies. Sensei expands the t-test and models the cell abundances using a beta-binomial distribution. We evaluate the mathematical accuracy of Sensei and provide practical guidelines on over 20 cell types in over 30 cancer types based on knowledge acquired from the cancer cell atlas (TCGA) and prior single-cell studies. We provide a web application to enable user-friendly study design via https://kchen-lab.github.io/sensei/table_beta.html.

## 2 Background

Cellular composition varies across different tissues and organs of the human body [1]. Cell type abundance is highly dynamic and varies across physiological and pathological states, including oncogenesis [2–4]. However, these changes in cell composition may be subtle and their detection requires the use of single-cell technologies coupled with accurate analytical pipelines allowing the enumeration of cell populations-of-interest with adequate specificity, especially for rare cell types [5]. Ascertainment of these changes is critical to understand the complexity of human diseases. For instance, the human immune system requires constant trafficking of different cell types to disease sites to mount innate and acquired immune responses [3, 4, 6]. In addition, the immune system has resident cells present in almost all organs [7, 8]. Observing temporal dynamics within the immune cell compartment is critical to understand processes such as autoimmunity [6, 9, 10], susceptibility to infections [6, 8], and development of cancers [3, 4, 6]. Changes in the abundance of specific immune cell types within the tumor microenvironment (TME) over time reflect the evolution of cancer across the successive stages of premalignancy, invasion, local recurrence and distant metastatic spread [5, 11, 12]. Differences in TME composition are also reflective on different subtypes of tumors associated with different coevolving immune responses, thus reflecting two of the hallmarks of cancer: evasion of immune detection, and tumor promoting-inflammation [13]. Therefore, these pieces of information are critical to understand the role of the immune system during cancer evolution and metastasis and also to develop immune interception strategies for both cancer prevention and treatment [2].

For example, the intestinal mucosa is populated by intra-epithelial lymphocytes and mucosa associated lymphoid tissue. Proportions of T cells may vary in mucosa specimens obtained from healthy individuals at average-risk for colon cancer development (general population) compared to individuals at high-risk as a consequence of genetic predisposition due to an inherited condition such as Lynch syndrome. Lynch syndrome is the most frequent hereditary syndrome predisposing for the development of colorectal cancer and is secondary to the presence of germline mutations in one of the DNA mismatch-repair (MMR) genes. The deficiency of this mechanism leads to the accumulation of hundreds of point mutations and insertion-deletion loops (indels) that generate hypermutant neoplastic lesions [14]. These mutations constitute antigenic peptides (also known as neoantigens) that are recognized by the immune system, thus leading to an activation of different immune cell populations. Therefore, studying changes in immune cell proportions at single-cell resolution could help understand the immune response triggered at the intestinal level, thus helping to envision strategies to enhance it to prevent cancer or to decrease it to treat conditions such as inflammatory bowel disease [15]. This type of study would require the use of multi-color flow cytometry [16] and also intersects with microbiome [17] datasets, but it can be now accomplished with much higher accuracy due to the rise of single-cell RNA-sequencing (scRNA-seq) and single-cell ATAC (assay for transposase-accessible chromatin) sequencing (scATAC-seq) [18, 19]. To observe and confirm cell type differences, samples from multiple research participants will need to be collected and sequenced; thus, accurately estimating the adequate sample size is critical for the feasibility and success of these type of studies due to the current high cost of these technologies. On the other hand, an insufficient number of samples can lead to a false-negative result [20].

Various sources of variability can complicate the ascertainment of cell type abundance. Sample preparation and single-cell sequencing reactions can introduce undesirable technical biases and variations [21]. For example, cell types that are hard to harvest intact such as neurons and adipocytes may be disproportionately underrepresented. As for single-cell profiling, scRNA-seq can introduce dropouts of lowly expressed genes, low total gene counts per cell, and high bias for 3’ coverage [5], while scATAC-seq can be confounded by sampling efficiency resulting in a highly sparse profiling [22]. Furthermore, mass cytometry brings its own challenges as it is susceptible to oxidization and signal spillover [23]. All of these factors often lead to uncertainty in cell typing and, therefore, need to be properly accounted for sample size estimation before the experiments are performed. Moreover, selection of the type of platforms relies on the number of cells that can be assayed, ranging vastly from 100 to 10,000 [5] and the fact that in many occasions few cells remain after performing quality control. In general, a limited number of cells leads to underrepresentation of cell types and drift in their proportions. Therefore, a method that considers these factors is urgently needed.

However, it is challenging to model the effects of these factors in a mathematical model. Several approaches have utilized statistical models to estimate the number of cells that are required for a single-cell study. “Howmanycells” (https://satijalab.org/howmanycells). It uses negative binomial distributions to estimate how many cells assayed in total ensure sufficient representation of a given cell type, assuming that the number of cells in different cell types are mutually independent. However, if the proportion of one cell type rises, the proportions of other cell types must fall. Accordingly, SCOPIT [24] uses Dirichlet-multinomial distribution to add negative correlations between cell types. Nevertheless, the authors of SCOPIT have verified that calculations based on the independence assumption are very similar to that of SCOPIT, only off by a maximum of one cell [24]. Further improvement in modeling is possible, but it will likely result in non-analytical solutions. Also, validating the accuracy of more sophisticated models will be unrealistic, as it requires datasets providing impractical and, most of the time, unfeasible numbers of technical replicates.

Most importantly, those previous approaches were designed to estimate the number of cells in a single biological sample, but not to estimate the number of biological samples that are required to ascertain changes in cell type abundance across biological conditions, a very different goal. For biological sample size estimation, the legacy sample size estimation approach for the t-test (Methods) does not factor in the variance introduced by insufficient number of cells. Thus, the estimation can be over-optimistic, especially for rare cell types.

Here, we present a new approach, Sensei, to provide accurate estimation of the sample size (or, equivalently, statistical power or false negative rate) for a variety of single-cell studies. Sensei takes into consideration both the number of samples and the number of cells within a unified mathematical framework and accounts for the abovementioned variabilities. We validate the accuracy of Sensei using multiple datasets and demonstrate Sensei’s utility in a wide range of study settings that can impact broadly on both cancer prevention and treatment. We have also developed an online web application making Sensei accessible for clinical and basic science researchers during a study design.

## 3 Results

### 3.1 Sensei

The framework of Sensei to model a controlled clinical study is illustrated in Fig. 1. The study design includes a control group and a case group of participants of certain sizes (Fig. 1a). The proportion of a cell type, T cell as an example hereafter, in a specific tissue varies among participants. While a level of difference is expected between the means of the T cell proportions in the two groups, within-group variances blur it, thus making statistical test necessary for ascertainment. Because a proportion falls between 0 and 1, Sensei uses a beta distribution to model the true proportion of T cells in each group, which parametrizes difference between groups and variance among participants within each group (Fig. 1b). For studies involving matched-pairs of specimens, e.g., autologous samples from one group of participants, additional statistical power can be acquired from modeling positive correlation of proportions of each cell type between pairs of samples.

**Fig. 1.**
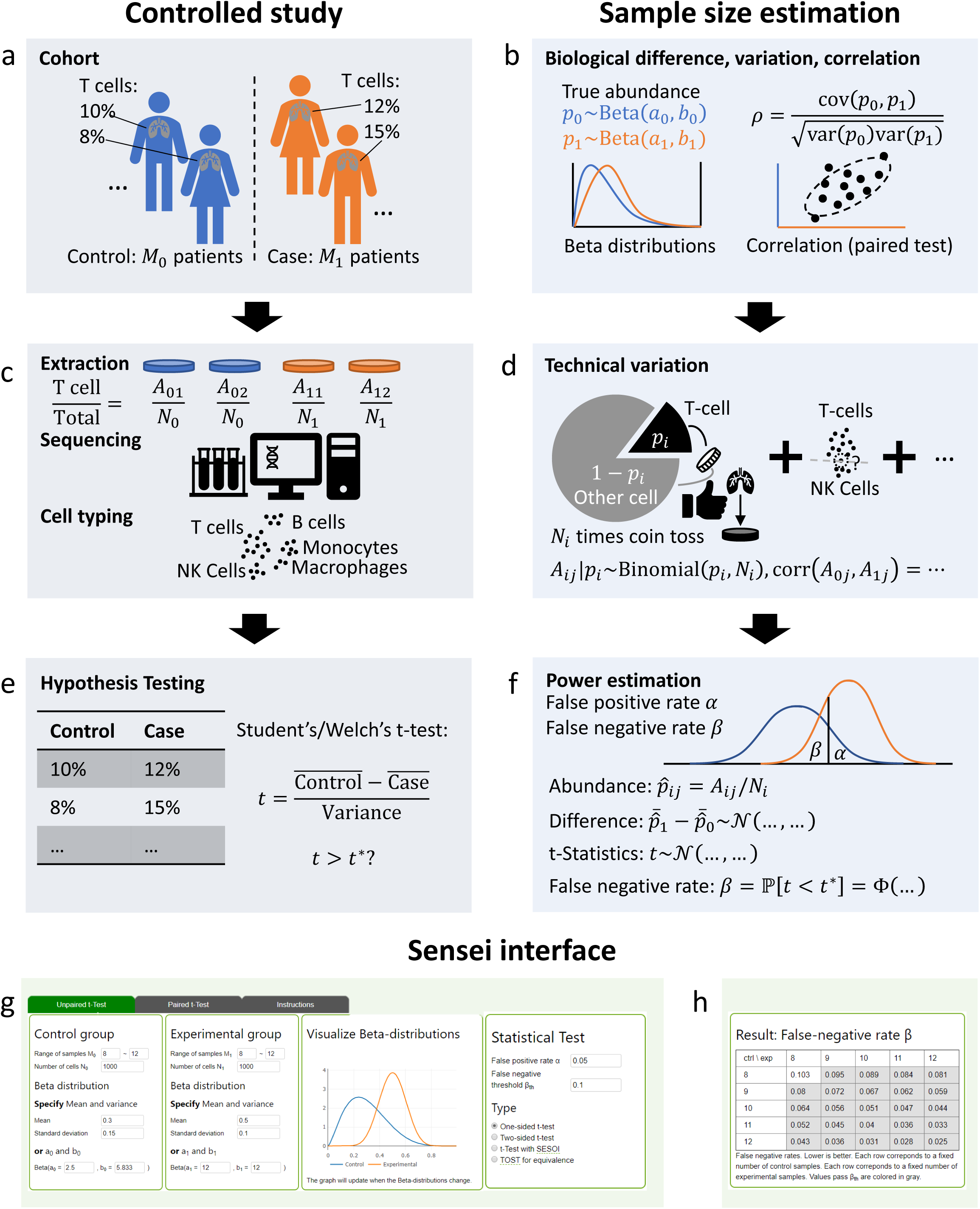
Framework of Sensei. **a**-**f** show side-by-side the way Sensei (right) models a controlled clinical study (left). **a** A controlled study involves a control group and a case group for ascertaining the difference in the proportions of T cells between the two groups. **b** Sensei models the true biological between-group difference and within-group variance using beta distributions. Correlation is also modeled for matched pairs study design. **c** A biopsy is extracted from each participant and assayed by a single-cell technology. Cell types are identified *in silico.* **d** Sensei models technical variations introduced by limited cell number using a binomial distribution (with other technical variations already accounted for in **b**). **e** The t-test is performed to identify statistically significant differences. **f** Sensei infers the distribution of the t-statistics and calculate the false negative (type II error) rates. **g** A sample input for Sensei. Required are sample sizes, cell numbers, estimated proportions of the cell type and false positive rate (type I error) rate for t-test. **h** A sample output of Sensei, corresponding to **g**. Tabulated are false negative rates for each feasible sample size.

From each participant, a biopsy of a tissue of interest is extracted, dissociated, and assayed using one of the single-cell profiling protocols. The single-cell profile is analyzed *in silico* and the cells are clustered and classified into cell types (Fig. 1c). Two types of technical variations are introduced in this step. Firstly, a major source of variation is limited cell number, especially for rare cell types, which reduces the statistical power of a study. To model it, we assume that profiled cells are chosen randomly from the population, i.e., all cells in the tissue of interest, which is consistent with SCOPIT [24] and “Howmancells”. Because the total number of cells in the tissue (population) is typically larger than that assayed in a single-cell experiment by several orders of magnitude, the number of sampled cells from a specific cell type would closely follow a binomial distribution, given its true proportion in the population (Fig. 1d). Secondly, sample preparation, sequencing and data analysis also raise uncertainty, which is highly complex and may not be modeled analytically. Precise modeling would require exhaustive quantification of a specific protocol, which is not readily available. Thus, we factor such variances in the beta distributions (Fig. 1b) mentioned above, which is consistent with the empirical understanding in the field [25]. The conjugacy of beta distribution and binomial distribution facilitates such modeling, allowing for efficient computation. Also factored in is the correlation between paired samples, if applicable.

After cell types are identified, assuming that the distributions of the proportions are approximately normally distributed, the t-test, one of the most widely used statistical tests [26, 27], can be applied to ascertain the between-group difference. Indeed, the observed skewness and kurtosis of cell type proportions validates the assumption of normality (Supplementary Text) [28] and justifies the use of the t-test. The t-statistics is calculated and compared with a critical value corresponds to a significance level (also referred to as false positive rate and type I error rate, 0.05 and 0.01 being the typical choice) (Fig. 1e). Sensei estimates the false negative (type II error) rate by inferring the distribution of the t-statistics and calculates the probability of it failing to reach the critical value (Fig. 1f). The correlation of samples in the paired test (Fig. 1b) is also accounted for.

Sensei is implemented as a web application powered by JavaScript, and as a Python package. Required as input are the sample sizes, cell numbers, estimated cell proportions, false positive rate and the type of t-test (Fig. 1g). Output is a table of false negative rates for various sample sizes for researchers to identify feasible study designs (Fig. 1h). An example for performing a paired t-test is shown in Supplementary Figure 1. Mathematical modeling is detailed in Methods.

### 3.2 Validation of Sensei

Because Sensei’s analytical solution includes necessary approximations (Methods, Equation 7 and Equation 10), we performed a simulation experiment to validate that Sensei accurately estimates the sample size for ideal beta-binomial distributions. We simulated 10,000 datasets using the beta-binomial model that Sensei aims to approximate (Methods). We set sample sizes *M*_0_, *M*_1_ = 5∼12, cell numbers per sample *N*_0_, *N*_1_ = 1,000, mean proportions *μ*_0_ = 0.03, *μ*_1_ = 0.05, and variances *σ*_0_ = 0.015, *σ*_1_ = 0.01 for control and case samples, respectively. We performed a one-sided unpaired t-test with a significance level of *α* = 0.05 on each dataset and counted the number of negative results to determine the false negative rate. We then used Sensei to estimate the false negative rates with the same parameters. For comparison, we also applied on the same data the legacy t-test approach, which makes predictions assuming a normal distribution instead of the beta-binomial distribution. As shown in Fig. 2a, the estimation error of Sensei against the simulated ground-truth (7.9% on average) is much smaller from that of the legacy approach (38.2%). The latter tends to be over-optimistic, because it does not account for insufficiency in cell number, which has relatively large effects on such a rare cell type.

**Fig. 2.**
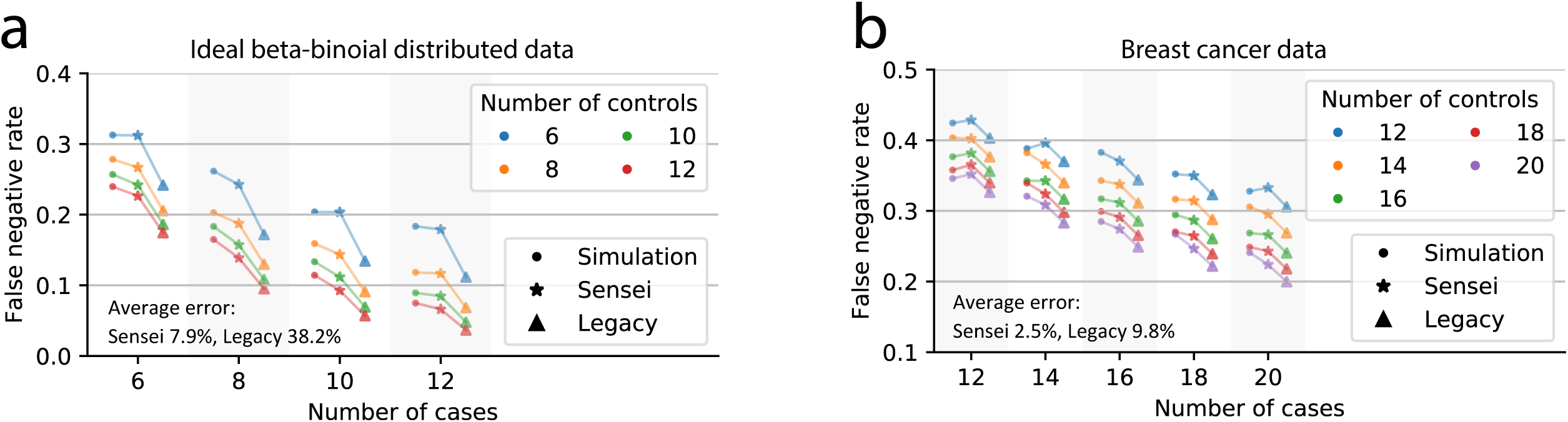
Results of simulation studies. **a** Comparison of false negative rate (y-axis) known from simulation against those estimated by Sensei and by the legacy approach, using datasets sampled from a beta-binomial distribution. Number of samples in the case group is indicated on the x-axis and in the control group by different colors. Markers correspond to result from different approaches. The average error is the mean absolute relative difference between the estimation and the simulation. **b** Comparison of false negative rates calculated by Sensei and the legacy approach, with those generated by simulation on the proportions of T cells in tumor and juxtatumoral samples in a breast cancer study.

Because real tissue data may not follow exactly the assumed distributions in the simulation, to further assess the accuracy of Sensei, we evaluated it on a breast cancer dataset, which contains 144 tumor samples, and 46 juxta-tumoral samples [29]. The proportions of T cells *p*_*ij*_ are available as ground truth for each sample, with an average of 56% in the tumor samples and 42% in the juxtatumoral samples (p-value = 6.6 × 10^−6^, two-sided t-test). We considered the tumor samples as the case group and the juxta-tumoral samples as the control group and assumed that a study plans to involve 12 to 20 participants per group to ascertain a change in T cell abundance. For each combination of sample sizes of both groups, we obtained the estimation from Sensei and the legacy approach using a simulated dataset generated according to the original data (Fig. 2b, Methods). A very high degree of consistency can be observed between the “Sensei” and the “Simulation” results (Fig. 2b). For 100 cells per sample, Sensei halved the average error of the “Legacy” approach (2.5% versus 6.6%). Because T cell is relatively abundant (Supplementary Figure 2), the improvement shrinks when more cells are collected (4.8% versus 5.7% for 384 cells and 3.8% versus 4.1% for 1000 cells). The improvement is expected to be larger for rare cell types. The result further validated the accuracy of Sensei in assessing immune cell abundance in breast cancer samples, which does not strictly follow the assumed distributions (Supplementary Figure 2).

With Sensei being validated, we comprehensively examined datasets from current large-scale cancer genomic studies that have over 30 cancer samples [2]. We applied Sensei to estimate how many samples are required to detect compositional changes in over 20 cell types in a particular cancer type. Our results can be utilized as a guideline for designing preclinical studies and clinical trials in a variety of settings.

### 3.3 Testing cell type abundance difference in unpaired cancer samples

Changes in tumor clonal fractions have been widely used to track cancer evolution dynamics [30–32]. As important are changes in immune cell abundance in the TME [33]. In many studies, case and control samples are collected from different groups of patients. Thorsson et al. [2] deconvolved bulk RNA-seq data from TCGA data (Supplementary Figure 3) using CIBERSORT and obtained the proportions of 22 immune cell types in 11,373 samples. The immune cells can be further grouped into 6 major types (T cells, B cells, NK cells, Macrophages, Dendritic cells, Mast cells). We obtained the sample mean, standard deviation, and confidence intervals of the proportion of each cell type in each cancer type (Methods, Supplementary Figure 4-6). Based on these inputs, Sensei inferred the sample sizes for ascertaining the difference between normal tissues and primary tumors in each cancer type using a one-tailed unpaired t-test at a significance level of 0.05 with at least 80% power (Fig. 3a, Supplementary Figure 7a,b). Although Sensei has the ability of suggesting unequal number of cases and controls, we assumed sample sizes are equal for both groups without loss of generalizability. The result shows that a sample size of 20 in each group is adequate to ascertain the differences of many cell types in many cancer types using current single-cell technologies, including but not limited to T cells in KICH (kidney chromophobe), KIRC (kidney renal clear cell carcinoma), READ (rectum adenocarcinoma), and THCA (thyroid carcinoma), and B cells in COAD (colon adenocarcinoma), ESCA (esophageal carcinoma), KIRC, KIRP (kidney renal papillary cell carcinoma), LUAD (lung adenocarcinoma), LUSC (lung squamous cell carcinoma), and READ (Fig. 3a).

**Fig. 3.**
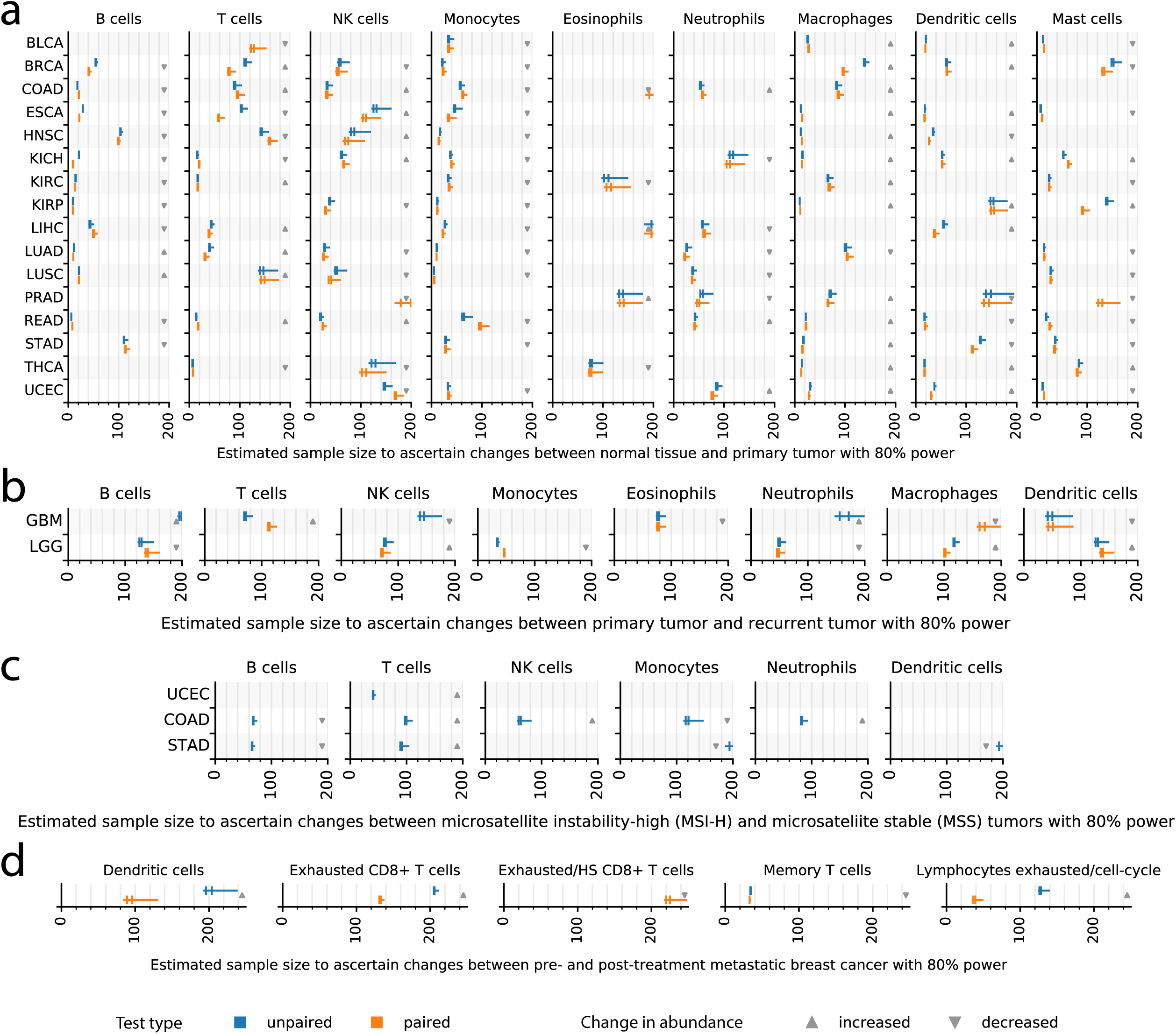
Sample size estimated by Sensei. **a** Estimated sample size for detecting statistically significant difference in normal tissue and primary tumor using an one-sided Welch’s t-test at a significance level of 0.05 with 80% power (the same below). Estimations for unpaired test and paired test are shown in blue and yellow, respectively. Estimations are for infinite (the legacy approach, left end of a whisker), 1,000 (left bar), 384 (right bar, may overlap with the left one), and 100 (right end of a whisker) cells. Fewer cells per sample would require more samples to ascertain an effect. The estimated sample size is for each of the two group in a controlled study, not jointly. For matched-pairs study, it is the same as the number of participants. Sample sizes larger than 200 are omitted. The direction of change in cell type abundance is shown by an arrow. An up arrow indicates a higher abundance in primary tumor compared with normal tissue, and vice versa. **b** Estimated sample size for detecting statistically significant difference in primary tumor and recurrent tumor for low grade glioma (LGG) and glioblastoma multiforme (GBM) patients. An up arrow indicates a higher abundance in recurrent tumor compared with primary tumor, and vice versa. **c** Estimated sample size for detecting statistically significant difference in each immune cell type between microsatellite instability-high (MSI-H) and microsatellite stable (MSS) tumor samples in uterine corpus endometrial carcinoma (UCEC), colon adenocarcinoma (COAD), and stomach adenocarcinoma (STAD). An up arrow indicates a higher abundance in MSI-H tumor compared with MSS tumor, and vice versa. **d** Estimated sample size for detecting statistically significant difference between pre- and post-treatment samples from metastatic melanoma patients. An up arrow indicates a higher abundance in post-treatment tumor compared with pre-treatment tumor, and vice versa.

Incidentally, a CyTOF study of LIHC (liver hepatocellular carcinoma) is available, involving 12 tumor samples and 7 normal tissue samples [34]. Sensei estimated a power of 75% for identifying an increase in regulatory T (Tregs) cells using the study sample size. Indeed, the study successfully detected an increase in Tregs at a statistically significance level between 0.05 and 0.01.

Similarly, we calculated the sample size needed for studying cancer progressions from primary tumors to recurrent tumors, since differences in the TME may indicate cancer metastasis and treatment resistance [35]. We have used a data set from a study assessing 13 samples of glioblastoma multiforme (GBM) and 18 of low grade glioma (LGG). Unlike tumor versus normal studies, the difference between recurrent and primary tumors is generally more subtle (Supplementary figure 4-6), and thus require more samples to ascertain (Fig. 3b, Supplementary Figure 7c,d). Our results show that compared with the primary tumors, a change in monocyte proportion in the recurrent tumors may be detected in LGG with a modest samples size of 34 (Fig. 3b). This is relevant, as previous studies have detected a significant decrease in monocyte proportions over malignant transformation of glioma [36]. A decrease in neutrophils proportion, which is known to be negatively correlated with glioma grade [37], also requires relatively modest sample sizes to detect. For GBM, Sensei predicts that a study design with 80% power needs at least 37 samples per group for dendritic cells, and more for other cells (Fig. 3b). A recent pivotal single-cell study finds 13 primary and 3 recurrent GBM samples are likely insufficient to ascertain changes in immune cell types [38]. Consistently, Sensei predicts a power of only 33% for dendritic cells, 9% for T cells, and even less for other cell types for such a setting. It should be noted that the data for recurrent tumors are limited in TCGA. Thus, more pilot experiments may be advised for designing related studies.

Cancer heterogeneity is driven by both genetics and epidemiology. Often performed are pan-cancer studies that categorize tumors based on shared genetic and/or epidemiological features [39, 40]. For example, patients with Lynch Syndrome or inflammatory bowel disease often develop colorectal cancers displaying high level microsatellite instability (MSI-H), while sporadic tumors more frequently display microsatellite stability (MSS). Molecular subtyping based on microsatellite instability is not only used in colorectal cancers, but also in other cancers such as endometrial and stomach tumors. Multiple clinical studies have shown that immune checkpoint-blockade therapy is more effective on MSI-H cancers, potentially because of a higher T cell infiltration rate compared to MSS cancers [41, 42]. To extrapolate those findings to a wider variety of cancer types, it is important to have a study design that can ensure the ascertainment of immune cell abundance.

As an example, we selected a set of MSI-H and MSS tumors samples in TCGA produced by Hause et. al [43]. In this dataset, MSI-H tumors comprise approximately 30% of uterine corpus endometrial carcinoma (UCEC), 20% of colon adenocarcinoma (COAD), 20% of stomach adenocarcinoma (STAD), and much lower in other cancer types (Supplementary Figure 8a). Using the cell type abundance deconvolved from the bulk RNA expression data [2] and the microsatellite instability labels obtained from genomic testing [43], we summarized the immune cell type abundance for the three cancer types (Supplementary Fig. 8b). For one-tailed unpaired Welch’s t-test at a significance level of 0.05 with 80% power, the sample sizes estimated by Sensei are summarized in Fig. 3c (and Supplementary Figure 7e,f). Testing for the higher proportion of activated memory CD4 T cells in MSI-H and MSS STAD requires the smallest sample sizes – 29, 26, 25 samples in each group with 100, 384, and 1,000 cells per sample, respectively (Supplementary Figure 7e).

The sample size estimated by the legacy t-test approach also reported 25 samples. It is no coincidence that it is the same as what Sensei estimated for 1,000 cells, because the legacy approach effectively assumes that there are infinite numbers of cells sequenced. Thus, the result suggests that a sample of 1,000 cells is enough in the sense that the variance introduced by cell number is neglectable compared with the within-group variance. On the other hand, having only 100 and 384 cells may compromise the statistical power. In fact, to detect the difference in NK cells in COAD, even 1,000 cells result in a sample size of 59 (Fig. 3c), which is higher than 56 obtained via the legacy approach.

Overall, there is a trade-off between the number of cells per sample and the sample size. A few other cell types, including CD8 T cells in UCEC (Supplementary Figure 7e) and activated NK cells in COAD (Supplementary Figure 7f), also require fewer than 40 samples per group. On the other hand, many cell types would require more than 200 samples per group (not shown in the figure). Those cases either correspond to a very small difference, or associate with very large variance (Supplementary Figure 8b). Caution must be exercised when designing experiments under those conditions.

### 3.4 Testing cell type abundance difference in paired cancer samples

Paired studies involve the use of autologous samples from the same patients and can reveal more pathologically relevant changes in cell type abundance. It is an ideal way for assessing differences in the TME between not only primary and metastasis/recurrent tumors, but also primary and adjacent normal samples [44, 45]. Cell type abundances are available for 717 patients from matched normal and primary tumor samples, and 36 patients from matched primary and recurrent tumor samples (Supplementary Figure 9a,b). For each cell type, we estimated correlations of the cell type abundances between paired samples in each cancer type (Supplementary figure 9c,d). We then calculated the sample sizes under the paired test settings using Sensei.

The result between paired normal and primary tumor samples is largely consistent with that of the unpaired test (Fig. 3a and Supplementary Figure 7a), while significantly smaller sample sizes are predicted for some cases. A salient example is the dendritic cells in LIHC (liver hepatocellular carcinoma), for which as low as 35 samples is needed, compared with 53 in the unpaired test. Similarly, difference in naïve B cells in BRCA can be revealed by 33 samples, instead of 47 in unpaired test (Supplementary Figure 7a). However, in some cell types, larger sample sizes are required because there are negative correlations between paired samples in the data (Supplementary Figure 9c,d). Those may be technical artifacts introduced by experimental and analytical variances, as we found no statistically significant negative correlations (see 95% confidence intervals for the correlations shown in Supplementary Figure 9c,d and explained in Methods). In practice, the expected correlation can always be adjusted based on accurate prior knowledge.

Similarly, we obtained result between paired primary and recurrent tumor samples for the lower grade glioma (LGG) and glioblastoma multiforme (GBM) (Fig. 3b and Supplementary Figure 7c,d), based on prior cell type abundances estimated from 14 LGG patients and 6 GBM patients. The result is largely consistent between the paired test and the unpaired test with some salient differences. For example, 30 samples per group are needed to ascertain the difference in activated NK cells in LGG, compared with 42 for the unpaired test (Supplementary Figure 7d). For GBM, the sample size needed for follicular helper T cells decreased to 32 from 116 (Supplementary Figure 7c). It is important to be able to examine at modest sample sizes these cell types, which have been reportedly linked to malignant transformation of glioma and associated with the prognosis [36].

Paired tests are often utilized to assess the safety and efficacy of a treatment. We examined a dataset containing 48 tumor samples from 32 metastatic melanoma patients treated with anti-PD-1 therapy, anti-CTLA4 therapy, and their combinations, among which paired pre- and post-treatment samples are available for 11 of the patients [46]. Across various immune cells, exhausted lymphocytes increase the most and memory T cells decrease the most in the abundances post treatment (Supplementary Figure 10a). We calculated the pre- and post-treatment correlations for each immune cell type. We found a strong correlation (0.79) of exhausted lymphocytes yet a weak correlation (0.06) of memory T cells (Supplementary Figure 10b). Based on these input parameters, Sensei infers that at least 35 and 124 samples are needed to ascertain increases in exhausted lymphocytes under paired and unpaired design, respectively (Fig. 3d). On the other hand, ascertaining decreases in memory T cells requires 33 and 34 samples, respectively. This exemplifies that researchers may benefit substantially from a matched-pairs study design when there is a clear positive correlation for a cell type of interest between paired samples (Supplementary Figure 10b), which can often be the case, as paired samples are most likely derived from the same genetic background and under similar physiological conditions.

### 3.5 Designing a precancer clinical trial for cancer prevention

Effective eradication of cancer relies on not only treatment but also prevention [47]. The AACR White Paper for cancer prevention [48] calls for acquisition of more longitudinal data from pre-cancer samples to facilitate the modeling of progression and regression of pre-cancerous lesions. Sensei can be of great use in designing such studies. We used Sensei to design a randomized, placebo-controlled clinical trial involving patients diagnosed with Lynch syndrome, as a continuation of a pilot study [15]. The objective of the study is to evaluate whether the experimental intervention leads to recruitment and/or activation of immune cells in colorectal mucosa. Participants will be randomized to receive placebo or the experimental drug for a total of 12 months. After the treatment period, colorectal tissue samples will be collected. The percentage of immune cells within the mucosa will be measured by scRNA-seq to determine whether there were significant differences between the mean percentage of immune cells in the experimental treatment arm versus that in the placebo. Based on in silico deconvolution of bulk RNA-seq data from untreated colorectal mucosa, we estimated that the immune cell population is approximately 18.6% at baseline, with a standard deviation of around 5%. We hypothesized that the population will increase by 10 percentage points to 28.6% in the experimental arm and the standard deviation will remain the same. Based on these pieces of information, Sensei estimated that for a one-sided t-test, 6 samples in each group is needed to yield a false negative rate *β* = 0.062 ≤ 0.1 if 1,000 cells are collected in each specimen (Table 1). Furthermore, if as many as 5,000 cells are collected in each specimen, then 5 samples in each group will be enough to achieve *β* = 0.1, i.e., 90% power.

**Table 1.** False negative rate for estimating T-cell abundance changes in colorectal mucosa

| Control | Experimental |  |  |  |  |  |  |
| --- | --- | --- | --- | --- | --- | --- | --- |
|  | 4 | 5 | 6 | 7 | 8 | 9 | 10 |
| 4 | 0.214 | 0.166 | 0.138 | 0.121 | 0.110 | 0.101 | 0.095 |
| 5 | 0.167 | 0.116 | 0.088 | 0.071 | 0.060 | 0.052 | 0.046 |
| 6 | 0.141 | 0.089 | 0.062 | 0.047 | 0.037 | 0.030 | 0.026 |
| 7 | 0.124 | 0.072 | 0.047 | 0.033 | 0.025 | 0.019 | 0.015 |
| 8 | 0.113 | 0.061 | 0.038 | 0.025 | 0.018 | 0.013 | 0.010 |
| 9 | 0.105 | 0.054 | 0.031 | 0.020 | 0.013 | 0.009 | 0.007 |
| 10 | 0.099 | 0.048 | 0.026 | 0.016 | 0.010 | 0.007 | 0.005 |

It should be noted that this experiment compares pre-cancerous tissues in a placebo-controlled study, which is different from comparing tumor with normal tissues in TCGA. Thus, the estimated sample size is different compared to that of COAD in Fig. 3a. Sensei can be broadly utilized in clinical trial design, as estimations of the prior parameters are often available from preclinical/pilot studies.

## 4 Discussion

Changes in cell type composition underlie cancer evolution and metastasis. Ascertainment of such changes is critical for understanding the coevolution of tumor and its microenvironment during carcinogenesis and responses to treatments. Single-cell assays have become viable ways to measure cell type proportions in each biological sample. However, of great needs is a reliable, comprehensive, easy-to-use tool, which estimates the number of samples required for ascertaining changes in cell type proportions between two group of participants. Unlike tools that are designed to estimate the number of cells for ascertaining cell type proportions in a single sample, Sensei is the first tool, to our best knowledge, designed to estimate the number of samples for ascertaining changes in cell type proportions, with the limited capacities on current single-cell platforms accounted for. Although necessary approximations are made, the estimation is accurate as indicated by its consistency with the result of computer simulation and of patient data from real experiments. Results from previous single-cell studies are also consistent in the ballpark with Sensei’s predictions [38]. Sensei runs in seconds on the front-end without requiring any connection to backend servers, providing versatile, secured utilities for researchers with limited resources.

The estimation of Sensei is based on several assumptions on existing single-cell profiling protocols. Firstly, the profiled cells are assumed to be chosen at random from the tissue of interest, which leads to the assumption of binomial distribution. An experimental validation to this assumption would require a large number of technical replicates profiled from the same biological sample, which is neither currently available, nor practically viable. Notably, the same assumption is adopted by SCOPIT [24] and “Howmanycells” and appears widely accepted. Choosing beta distribution conveniently models cell type proportion among participants and greatly facilitates efficient computation via beta-binomial conjugacy. Further, the beta distribution can be uniquely determined by a mean and a standard deviation, which are widely accessible from preclinical studies. Like the beta distribution, its multidimensional extension, Dirichlet distribution, has been used in similar contexts [24]. More realistic modeling is possible, should become available more prior knowledge about biological variances, technical noise, and experimental biases, although an analytical solution may not exist. The power estimation will then be based on sampling, which requires substantially more computational resources. These pieces of information may not become clear, until experimental protocols become standardized and large single-cell atlases are completed [48–50].

In rare cases where the assumptions are violated, researchers should be able to observe a large skew in the distribution in data analysis. In those cases, new single-cell-aware power estimation methods based on non-parametric Wilcoxon rank-sum test might be more advisable [27, 28]. Sensei also assumes that the ascertainment bias within individual studies is consistent and well-controlled across experiments, i.e. equally applied to all the study samples and there are no significant batch effects, or that the batch effects have been alleviated by other systematic approaches. If severe batch effects are expected samples, stratified sampling, stratified test [51], and corresponding power estimation methods [52] should be used.

The effect on the false-negative rate of the total number of cells in each sample *N*_*i*_ is generally minimal, when the number of cells is greater than 1,000. Only for rare cell types (<5% proportion) will further increasing *N*_*i*_ become necessary to ensure statistical power. Our model assumes that *N*_*i*_ is the same for all samples in group *i*, which is a reasonable simplification because the number of cells generated by an assay is usually consistent in a systematic study and that small differences in *N*_*i*_ have little effect on results. It should also be noted that the standard deviation *σ*_*i*_, an input of Sensei, does not explicitly delineate biological variance among participants from technological variance introduced by assays. That said, those variances often coexist, and can hardly be separated cleanly. Thus, it is pragmatic to use the total variance that are learnt from existing or preliminary studies. Sensei’s estimation is based on Welch’ version of t-test, which also handles two groups with different variances, in addition to the standard Student’s t-test. Overall, our current model allows for a closed-form representation of the statistical power, which is essential to a light-weight web-based application providing a fast and sufficiently accurate estimation of sample sizes.

For convenience of presentation, we showed results based on an equal number of samples in the case and the control group, even for unpaired test. That is not a limitation of Sensei. In practice, the number of samples are allowed to be different between the two groups for unpaired studies. It should be noted that decreasing the number of normal tissue samples usually has less pronounced effect on the statistical power. Generally, the group with less variance requires fewer samples. Researchers may use our online application to choose the best combination of sample sizes for the two groups.

Sensei contains an implementation of additional variants of the t-test, including the smallest effect size of interest test and equivalence test (Methods), to support different kinds of studies. We have shown that the t-test is appropriate for most cell types, based on the TCGA data. We have also shown that the correlations of cell type proportions between paired samples are positive for many cell types, which empowers the paired test. We also provide a guideline for setting the parameters including mean, variance, and correlation in Sensei.

We expect that Sensei, with rich information we summarized from various datasets including normal/tumor, primary/metastasis/recurrent tumor, and pre-/post-treatment data, will meet the demand of many projects that are being planned, such as those in the Human Tumor Atlas Network [53] Pre-Cancer Atlas [48, 50], and clinical trials. Similar single-cell studies are on the rise at present. For example, even for colorectal carcinoma, where a relatively large cohort of data have been collected, more samples for colorectal adenoma are still needed to study the recruitment of immune cells throughout the lesion to find interventions that intercept premalignancy and prevent cancer [47]. In turn, data collected from these projects will inform Sensei to provide more realistic estimate.

## 5 Conclusions

This study reports a user-friendly web application for estimating sample size and statistical power in studies that apply single-cell profiling technologies to compare cell composition across samples. Both the number of participants and the number of cells per sample are taken into consideration. With an emphasis on cancer evolution, our results provide a guideline for designing studies to ascertain changes in cell type abundance among normal/tumor, primary/metastasis/recurrent tumor, and pre-/post-treatment conditions. We expect that Sensei will have applications in different single-cell studies involving differential abundance analysis. The web application can be accessed at https://kchen-lab.github.io/sensei/table_beta.html [54].

## 6 Methods

### 6.1 Beta-binomial modeling of Sensei

We assume that the study design includes *M*_0_ and *M*_1_ participants in the control and case group, respectively. For each group,*N*_*i*_ (*i* = 0, 1) single cells are collected in each sample. The mean *μ*_*i*_ and the standard deviation *σ*_*i*_ (as a result of biological variation) represent the proportion of the cell type of interest in each group. The significant level (false positive rate, normally 0.05 or 0.01) *α* should be assigned based on the expectation of the study, to calculate the false negative rate *β*, or, equivalently, the statistical power (1 − *β*). The input parameters required to execute our method are summarized in Table 2.

**Table 2.** Required parameters

| Parameter | Notation |
| --- | --- |
| Number of samples under condition $i = 0$ (control), 1 (case) | $M_i$ |
| Number of cells in each sample | $N_i$ |
| Mean and standard deviation of proportions for each beta distribution | $\mu_i, \sigma_i$ |
| False positive rate | $\alpha$ |

We assume that in the tissue to be studied, the true proportion of the cell type of interest, is *p*_*i*_. For the *j*th participant in group *i*, we denote *A*_*ij*_ the total number of such cells which is a random variable and has the following conditional distribution,

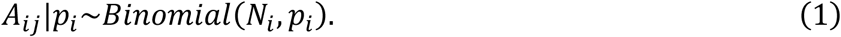

Because *p*_*i*_ is largely unknown in real cases, we model *p*_*i*_ using the conjugate prior of binomial distribution,

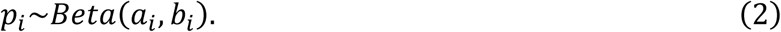

Therefore, the cell number *A*_*ij*_ have the beta-binomial distribution,

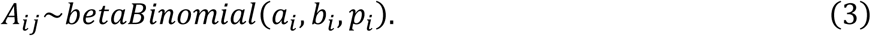

It is worth mentioning that beta-binomial distribution has been applied on modeling in compositional analysis [25, 55]. It is also a simplified version of Dirichlet-multinomial distribution used in sample size calculation [24, 56]. The *a*_*i*_ and *b*_*i*_ can be reparametrized from the user-defined mean and standard deviation *μ*_*i*_ and *σ*_*i*_. Formally,

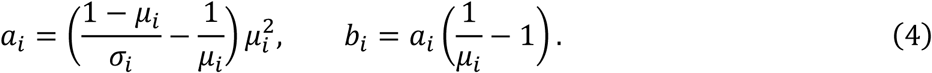

Practically, we require that the resulting *a*_*i*_ and *b*_*i*_ to be both greater than 1 to confine the beta distribution to be of unimodality. Using the properties of beta binomial distribution, we can get

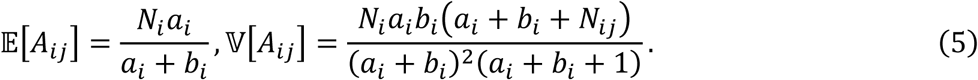

The corresponding cell type proportion is defined as 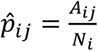 which follows a scaled beta binomial distribution. Thus,

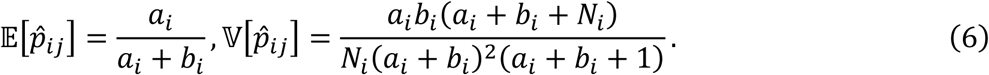

We now assume that the beta binomial distribution can be approximated by a normal distribution

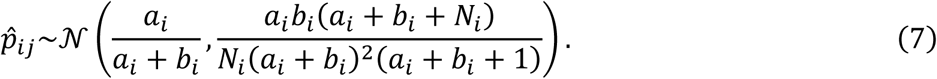

The approximation is justified by the fact that the L1 distance between the scaled beta-binomial distribution and Equation 7 is sufficiently small, especially for large *NN* and small *σσ* (Supplementary Figure 11a). We experimented on a few examples, for *μ* = 0.3, *σ* = 0.2, the underlying beta distribution is skewed to the left and deviates from a normal distribution. That results in a slightly unprecise, but still largely acceptable normal approximation (Supplementary Figure 11b). For *μ* = 0.5, *σ* = 0.1, the beta distribution itself is already close to a normal distribution, and the generated 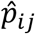 can be perfectly approximated by a normal distribution (Supplementary Figure 11b).

For a two-sided test, the null hypothesis is formulated as

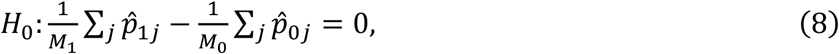

where 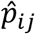 denotes the cell proportion in sample *j* from group *i*. For a one-sided test, the “=” is substituted by “<” or “>“. Thus, for a t-test allowing different variances in two samples [57], the t-value in Welch’s t-test follows a noncentral t-distribution, i.e.,

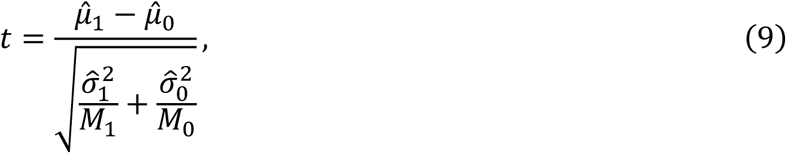

where the 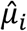 and 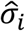 are sample mean and sample standard deviation of 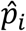, which are random variables. The distribution of t can be approximated by

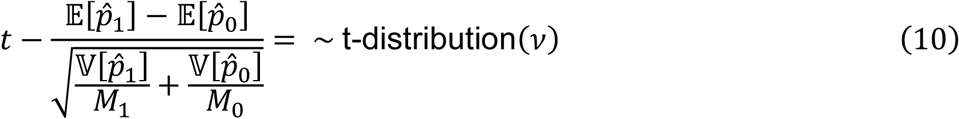

where the second term is a constant. The degree of freedom, *v*, is calculated as

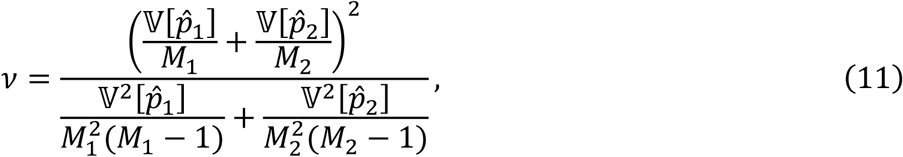

which degrades to (*M*_1_ + *M*_2_ − 2), the same as Student’s t-test, when 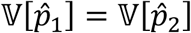 and *M*_1_ = *M*_2_ [57]. Thus, the false negative rate can be calculated as

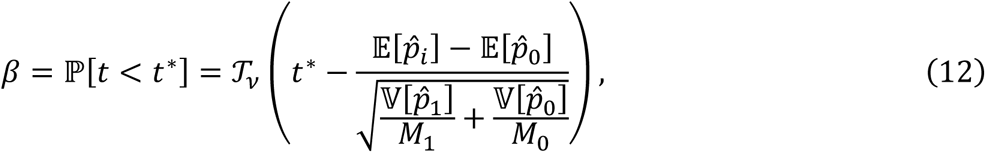

where 𝒯_*v*_, is the CDF of the Student’s t-distribution. 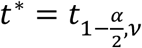, as 2ℙ[t ≥ t*] < *α*, for a two-sided test [58], or t* = t_1−*α,v*_ for a one-sided test.

### 6.2 Paired test

Paired samples are usually collected from normal and malignant tissues, or primary and recurrent/metastatic tumors. Longitudinal data from one patient, such as pre-treatment and post-treatment also form paired samples. In such cases, paired test can exploit the correlation between paired samples to improve the statistical power. Sensei has a functionality to help design studies with paired samples. In addition to the unpaired test, we naturally require sample size *M*_0_ and *M*_1_ to be the same (denoted as *MM*) and require one more parameter, *ρ* = corr(*p*_0_, *p*_1_), the correlation of the true proportions of cells between two conditions in the paired study. Note that cell number of cell type *A*_0_ and *A*_1_ are solely depend on *p*_0_ and *p*_1_, respectively. Thus, they are conditionally independent given *p*_0_ and *p*_1_. Consequently, we can use law of total covariance to derive

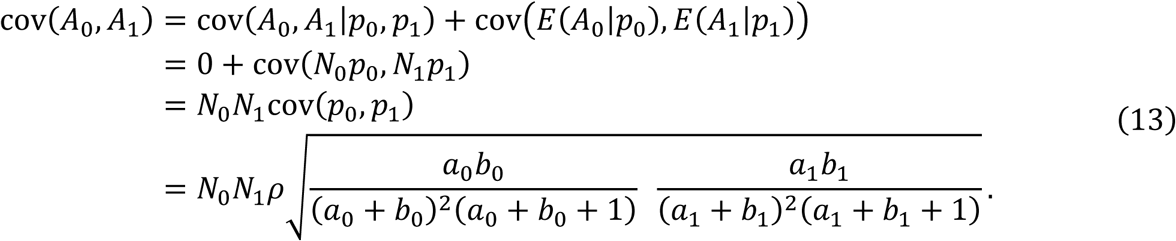

Thus, we have the distribution of the cell numbers

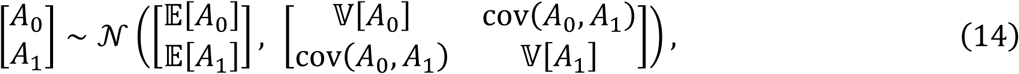

and proportions

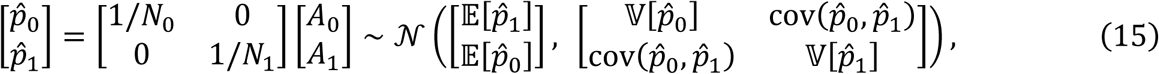

where 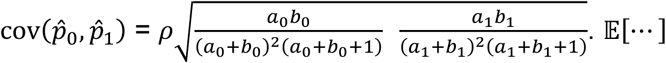 and 𝕍[…] remains the same as those in unpaired test. Note that 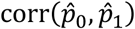 is in fact 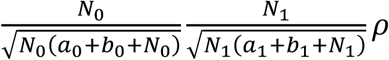, which approaches the same as *ρ* when numbers of cells, *N*_0_ and *N*_1_ are large. The difference between a pair of samples is

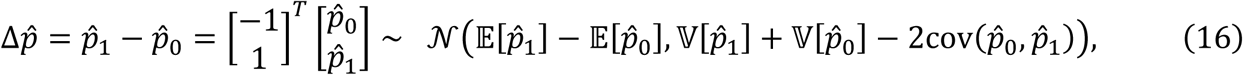

Thus, the paired t-statistics can be calculated as

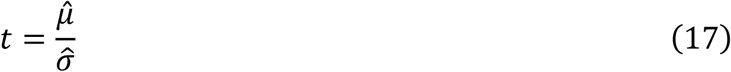

where 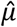 and 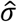 are sample mean and sample standard deviation of 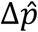. Thus, *t* satisfies

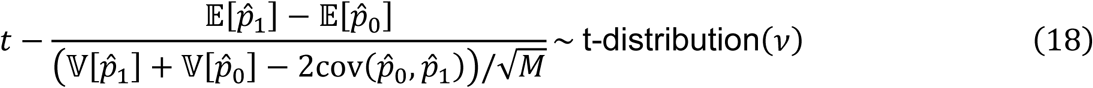

It can be observed that the t-statistics will be the same as the unpaired test when the covariance is zero, and even smaller should the covariance be negative. In other words, paired test needs a positive correlation to gain statistical power. Also note that paired t-test does not assume an equal variance. Finally, the false negative rate is

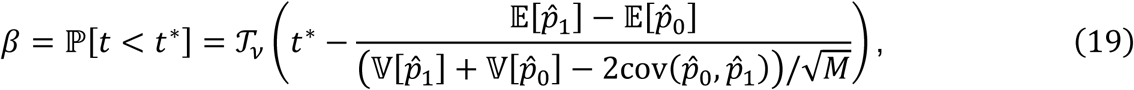

where t* = t_1−*α*/2,*v*_ for a two-sided test, or t * = t_1−*α,v*_ for a one-sided test, where *v* = *M* − 1.

### 6.3 Legacy Sample Size Estimation

We refer to the sample size estimated using the mean, variance, and correlation without the beta-binomial modeling in Equation 5 and Equation 13. Consequently, the effect of number of cells is not accounted for. It is effectively assuming an infinite number of cells.

### 6.4 Smallest Effect Size of Interest and two one-sided t-test for equivalence

Being Scientifically significant is usually different from being statistically different. For example, when enough samples are collected, even a 0.01% change in the proportion of a cell type can be statistically significant. However, the difference may be too small to induce any actual effect, and thus is rarely considered biologically interesting (i.e., not scientifically significant). Smallest effect size of interest (SESOI) is a way to set a threshold of scientific significance into statistical test [59]. Instead of performing t-test on the experimental group with the control group directly, it translates the control group by SESOI, the level to be considered biologically interesting, by adding or subtracting a constant from the control group. SESOI can also be used on the opposite side, to conclude that it is statistically significant, that the change in cell type abundance does not exceed the SESOI. We provide sample size estimation for t-test with SESOI in Sensei.

If two t-test with SESOI find that the different is statistically significantly within a range that is considered negligible in terms of biology, the proportion can be claimed to be effectively unchanged. This approach is formally called two one-sided t-test (TOST) for equivalence [59]. Sensei can also estimate the sample size for TOST.

### 6.5 Mean, variance, correlation, and their confidence intervals

The correlation and its confidence interval are obtained by standard ways [60], i.e., for cell type proportions in matched pairs {(*p*_0*i*_, *p*_1*i*_)}, *i* = 1 … n, the sample correlation coefficient and its (1 − *α*) confidence limits are

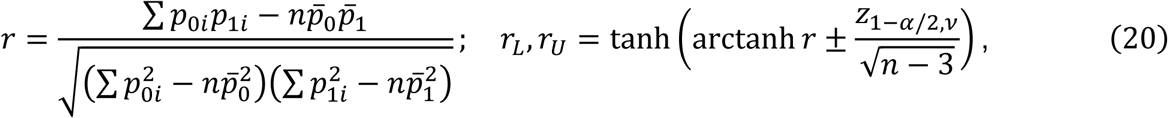

where *v* = *n* − 1. The sample mean 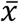, variance *s*, and their 95% confidence intervals 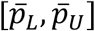 and [*s*_*L*_, *s*_*U*_] are obtained by standard methods for sample mean and sample standard deviation, i.e., for a group {*p*_*i*_}, *i* = 1 … n,

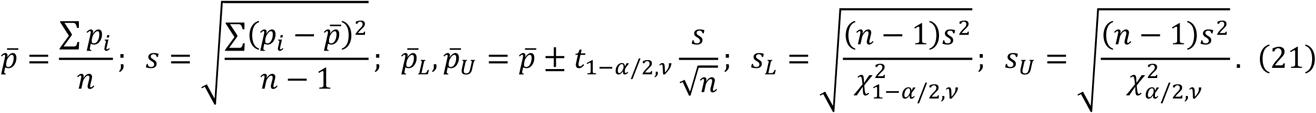

Sensei may use 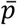 and *s* as input directly because they are the maximum likelihood estimates of parameters of a beta distribution. The confidence intervals may help evaluate the reliability of the prior knowledge. Note that the confidence limits may exceed [0, 1] in some cases, and we cut it to 0 or 1 in such cases. As a footnote, complementary log-log transform may be used to confine the limits, but it also skews the values and complicates interpretation. Bootstrap may also be used to construct the confidence interval.

### 6.6 Simulation study based on T cell abundance in breast cancer data

The breast cancer dataset contains 144 tumor samples, and 46 juxta-tumoral samples [29]. The proportions of T cells were available as ground truth for each sample, with an average of 56% in the tumor samples and 42% in the juxta-tumoral samples. We considered the tumor samples as the experimental group and the juxta-tumoral samples as the control group. Because the proportions of T-cells are significantly different (p-value = 6.6 × 10^−6^, two-sided t-test) between the two groups, we assume that true difference exists. We use the mean and standard deviation calculated as the input of Sensei. To validate Sensei’s accuracy, we randomly drew *M*_0_ and *M*_1_ samples respectively from the juxta-tumoral and tumor samples. If we were to perform single-cell assays on these samples, we would observe *A*_*ij*_ T cells in each sample, according to a binomial distribution parameterized by *N*_*i*_ and *p*_*ij*_ (*i* = 0,1). Binomial distribution is a reasonable assumption since a tissue sample often contains millions of cells, which is several orders of magnitudes higher than *N*_*i*_. We then perform a one-tailed unpaired t-test between the set of {*A*_0*i*_} and that of {*A*_1*j*_} at *α* = 0.05, and record a true positive when the test is positive, and a false negative otherwise. We estimate the false negative rate by repeating the above process 1,000 times for each combination of *M*_0_ and *M*_1_

## 7 Declarations

### 7.1 Ethics approval and consent to participate

Not applicable.

### 7.2 Consent for publication

Not applicable.

### 7.3 Availability of data and materials

The datasets analyzed during the current study are available with the original publications: Breast Cancer (https://doi.org/10.1016/j.cell.2019.03.005, Table S5) [29], Metastatic melanoma (https://www.ncbi.nlm.nih.gov/geo/query/acc.cgi?acc=GSE120575) [46], deconvolved TCGA (https://gdc.cancer.gov/about-data/publications/panimmune, Cellular Fraction Estimates) [2], and MSI-H and MSS labels (https://doi.org/10.1038/nm.4191, Supplementary Table 5) [43]. The web application, Python package, and code to reproduce the results are available at https://github.com/KChen-lab/sensei.

### 7.6 Authors’ contributions

All authors conceptualized the research. SL and JD conceived the statistical model. SL implemented the software and performed the experiments, with input from JW, JD, VM, YH, EV, and KC. All authors prepared and approved the final manuscript. EV and KC supervised the work.

### 7.8 Authors’ information (optional)

## 7.7 Acknowledgements

We thank Alex Davis and Nicholas Navin for their comments.

## 10 Additional Files

Additional File 1: Samples available for relative abundance of immune cells in the deconvolved TCGA dataset, related to Fig. 3a and Supplementary Figure 3.

Additional File 2: Sample mean, standard deviation, and their 95% confidence intervals for relative abundance of each cell type in additional - new primary samples, related to Fig. 3a and

Supplementary Figure 4-6. Cell types without enough information to make such inference are left blank. The same applies to other files below.

Additional File 3: Sample mean, standard deviation, and their 95% confidence intervals for relative abundance of each cell type in additional metastatic samples, related to Fig. 3a and Supplementary Figure 4-6.

Additional File 4: Sample mean, standard deviation, and their 95% confidence intervals for relative abundance of each cell type in metastatic samples, related to Fig. 3a and Supplementary Figure 4-6.

Additional File 5: Sample mean, standard deviation, and their 95% confidence intervals for relative abundance of each cell type in primary blood derived cancer - Peripheral Blood sample, related to Fig. 3a and Supplementary Figure 4-6.

Additional File 6: Sample mean, standard deviation, and their 95% confidence intervals for relative abundance of each cell type in primary solid tumor samples, related to Fig. 3a and Supplementary Figure 4-6.

Additional File 7: Sample mean, standard deviation, and their 95% confidence intervals for relative abundance of each cell type in recurrent solid tumor samples, related to Fig. 3a and Supplementary Figure 4-6.

Additional File 8: Sample mean, standard deviation, and their 95% confidence intervals for relative abundance of each cell type in solid tissue normal samples, related to Fig. 3a and Supplementary Figure 4-6.

Additional File 9: Sample mean, standard deviation, and their 95% confidence intervals for relative abundance of each cell type in the metastatic melanoma cancer dataset, related to Fig. 3d and Supplementary Figure 10.

