## Supplementary Figures for "Sensei: How many samples to tell evolution in single-cell studies?"

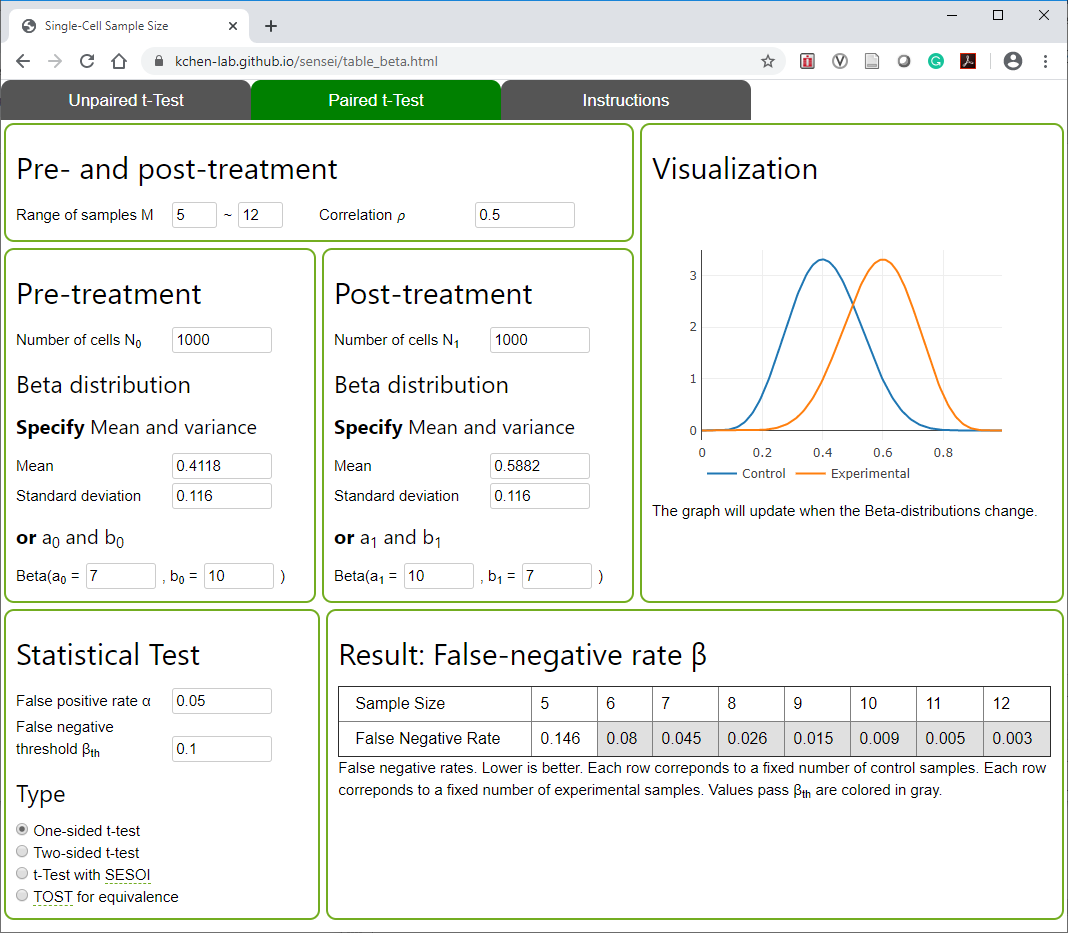

**Supplementary Figure 1**. Screenshot of Sensei for paired samples.

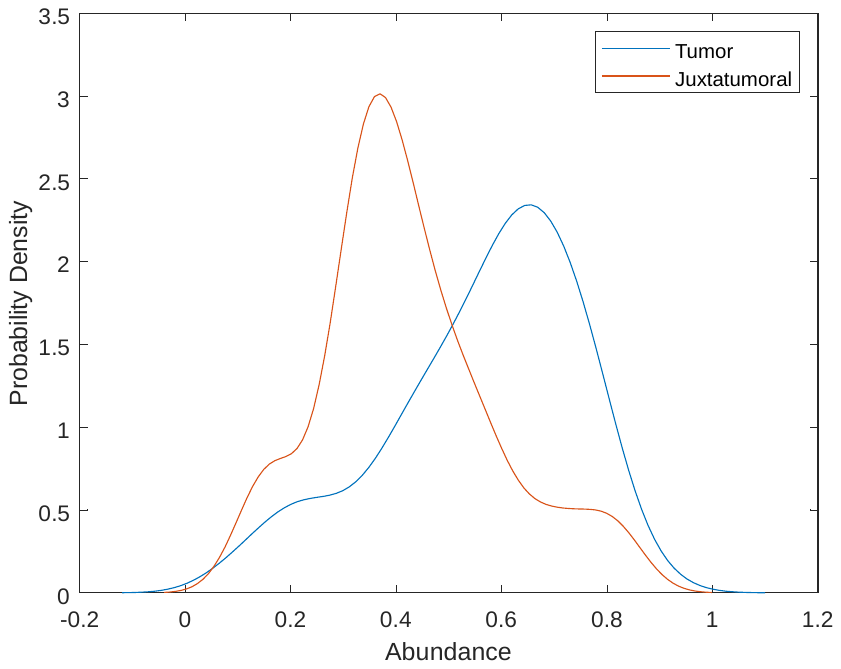

**Supplementary Figure 2.** The empirical distribution of the T-cell abundance in tumor group and juxtatumoral group, respectively.

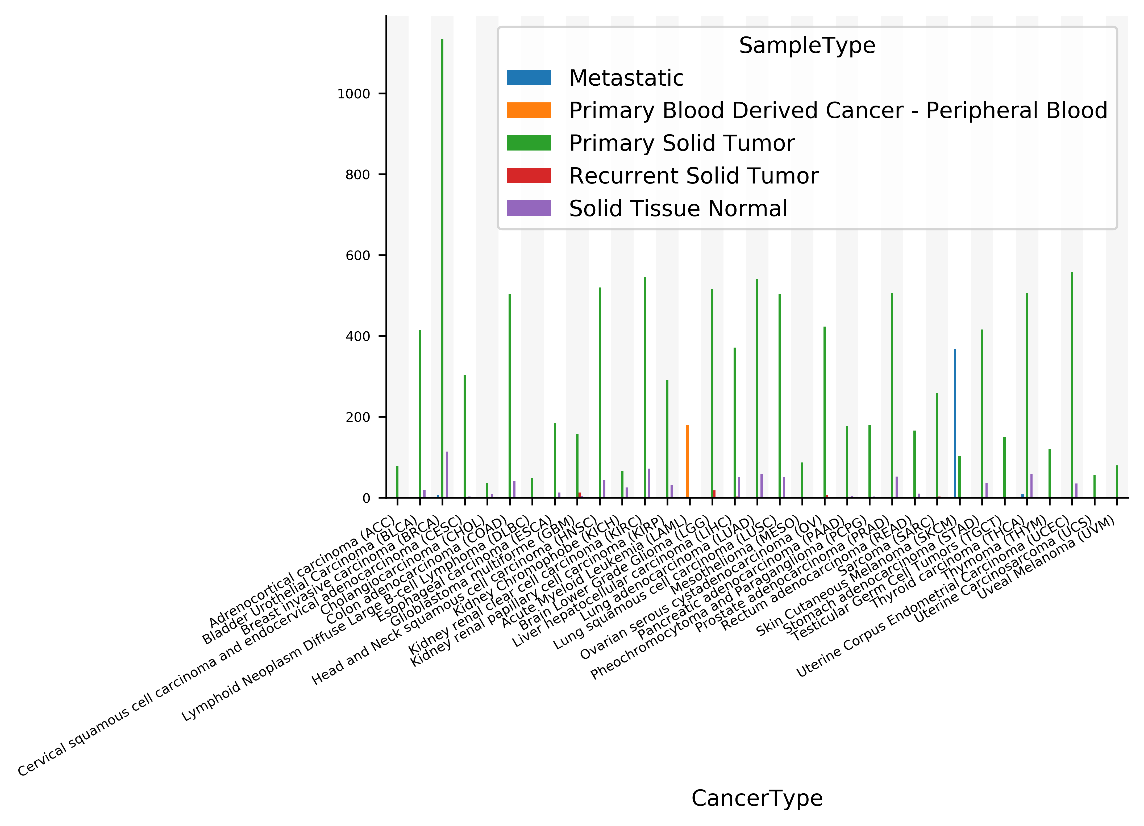

**Supplementary Figure 3.** Number of samples available in TCGA for each sample type in each cancer type.

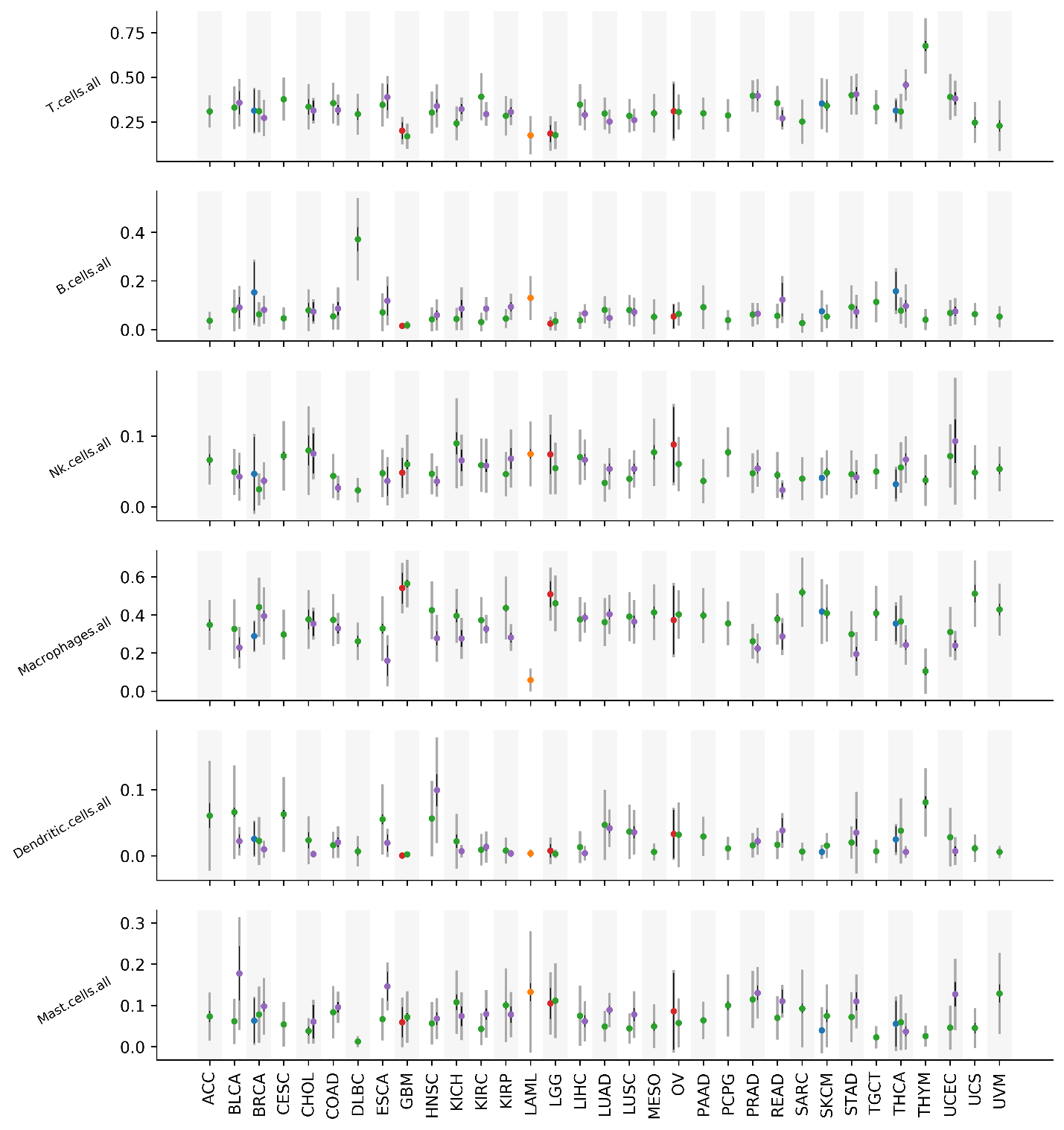

**Supplementary Figure 4.** Mean (dot), 95% confidence interval of mean (thin black line), and standard deviation (wide gray line) of the abundance of each major immune cell types in the deconvolved TCGA data. Color correspond to sample type with the same scheme in Supplementary Figure 3.

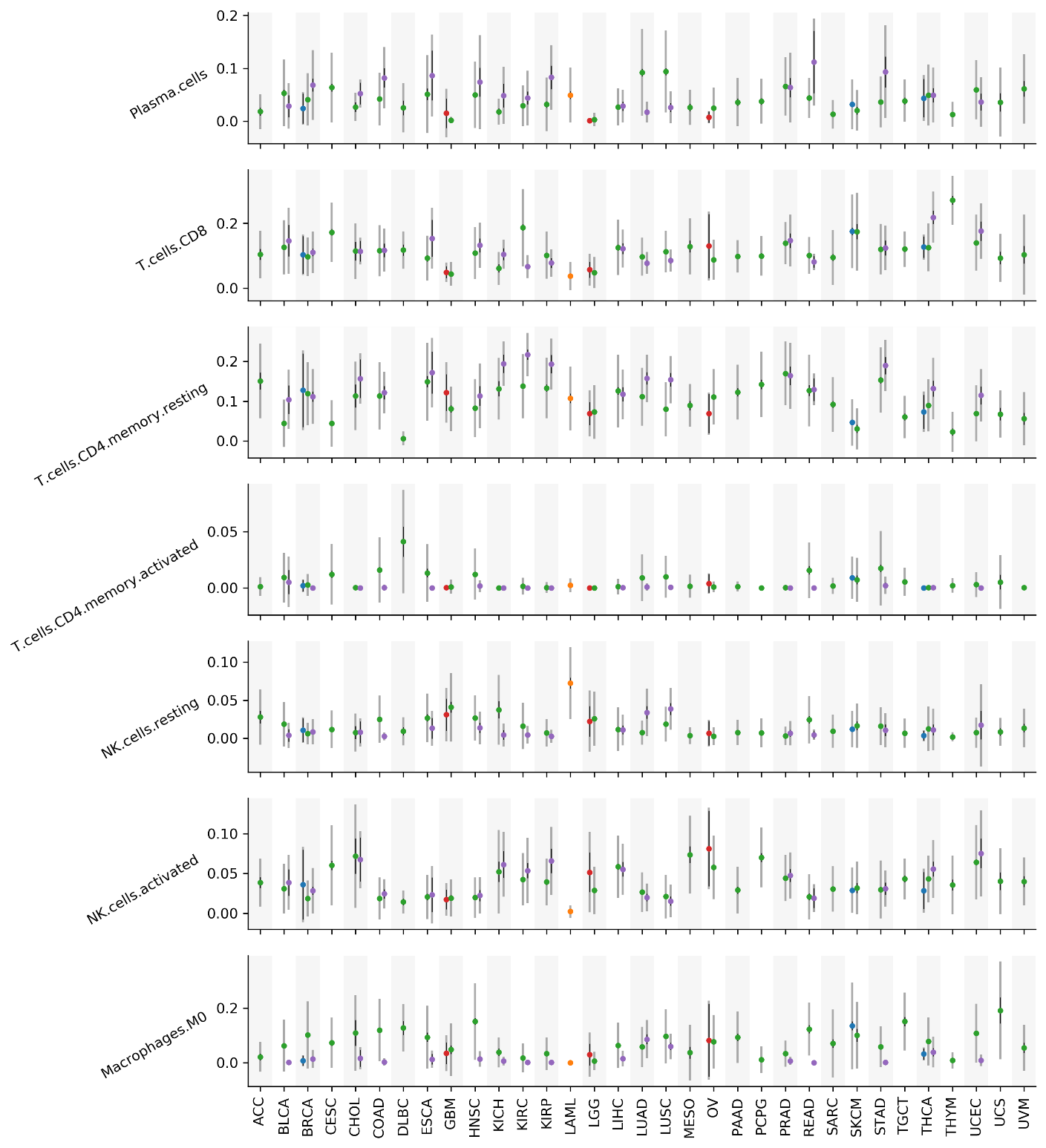

**Supplementary Figure 5.** Mean, 95% confidence interval of mean, and standard deviation the abundance for selected immune cell types in the deconvolved TCGA data. Scheme is the same as Supplementary Figure 4. Remaining cell types are shown in Supplementary Figure 6.

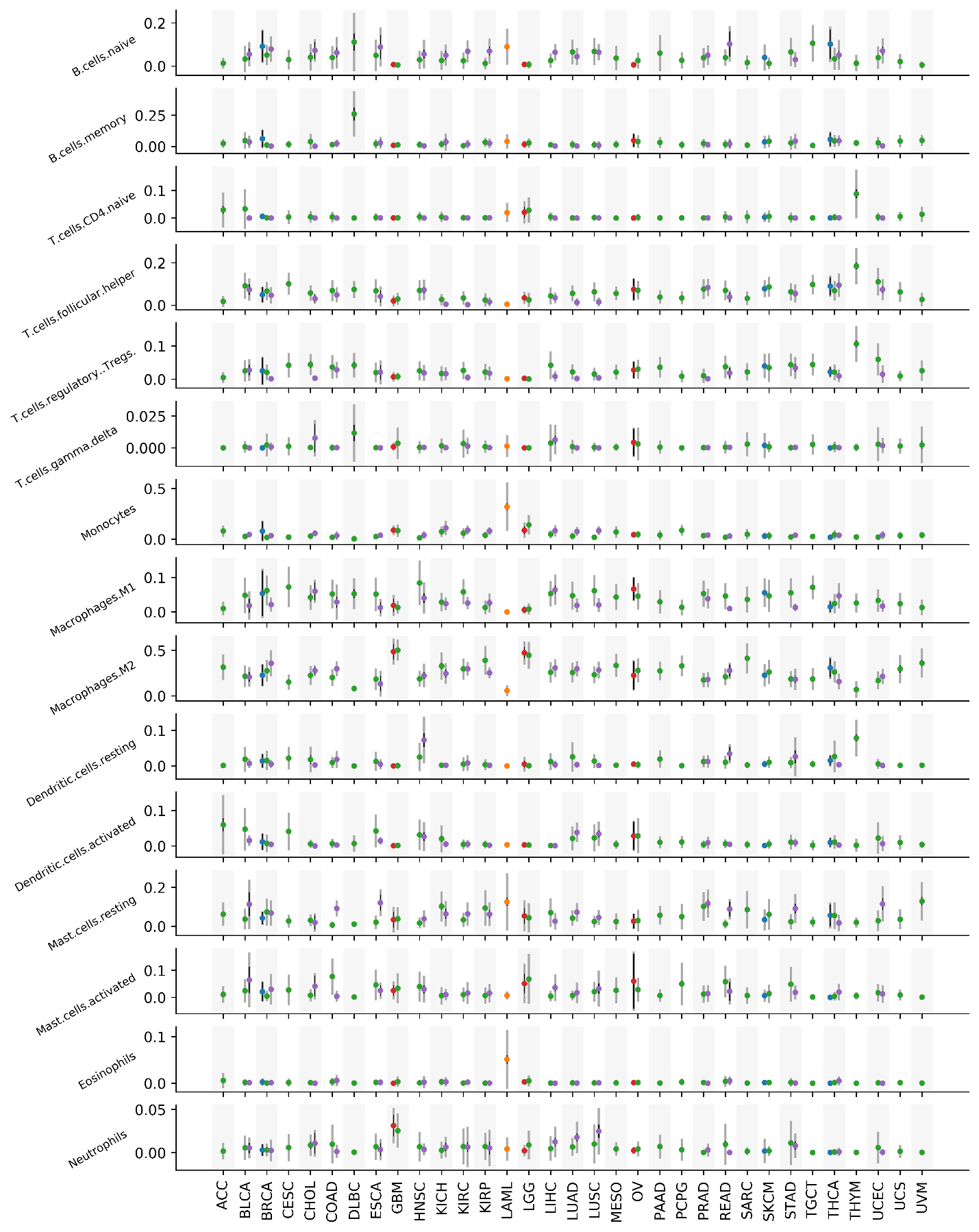

**Supplementary Figure 6.** Mean, 95% confidence interval of mean, and standard deviation the abundance for remaining immune cell types in the deconvolved TCGA data. Scheme is the same as Supplementary Figure 4 and 5. Other cell types are shown in Supplementary Figure 5.

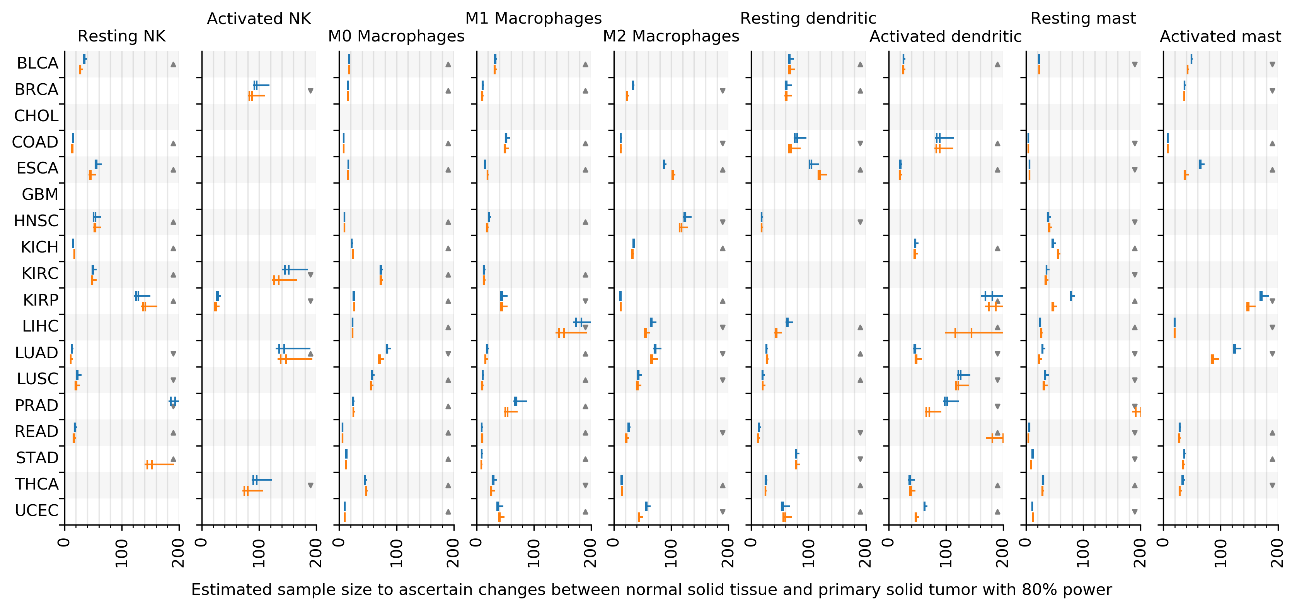

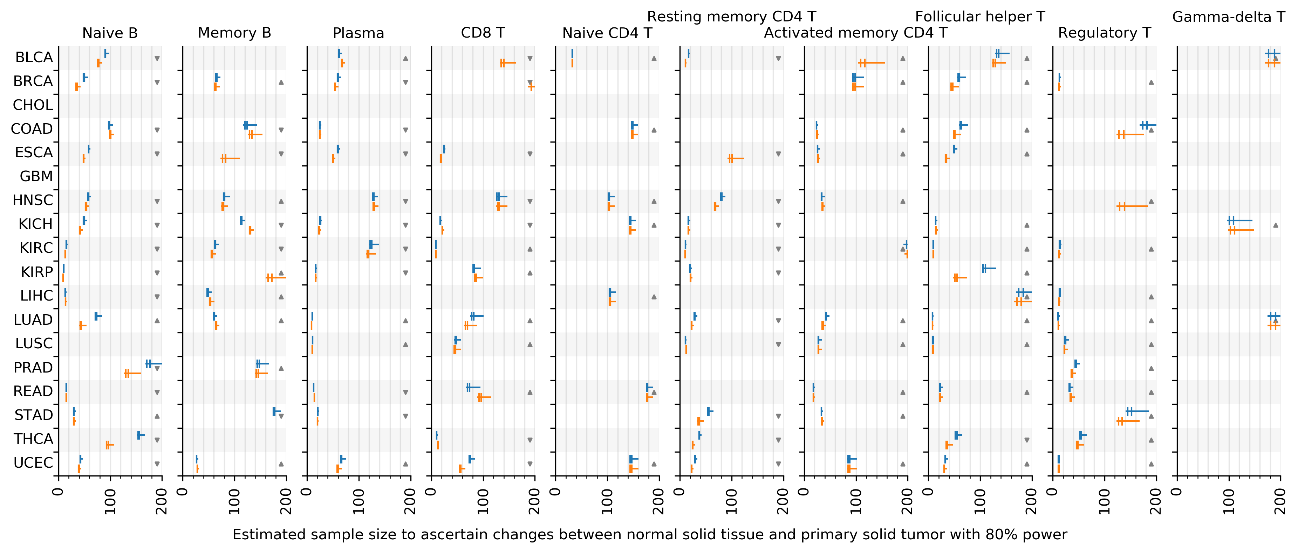

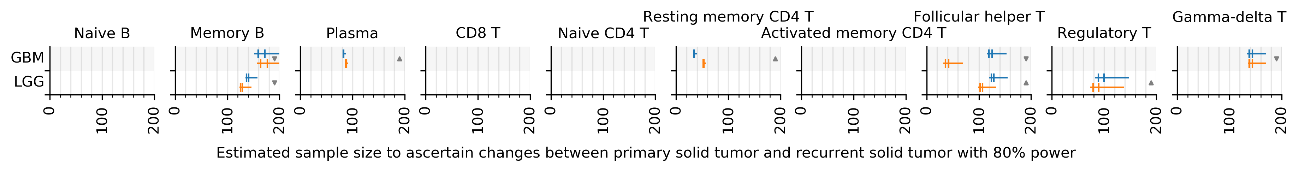

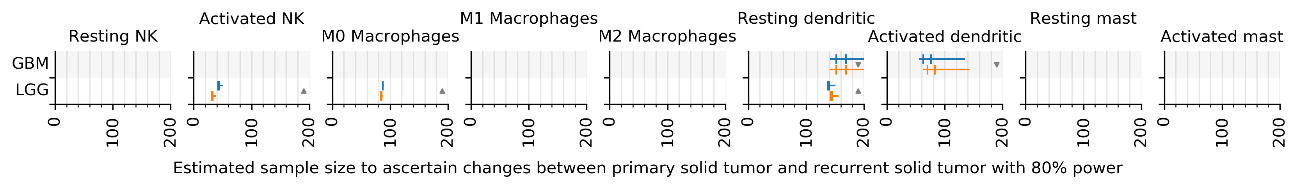

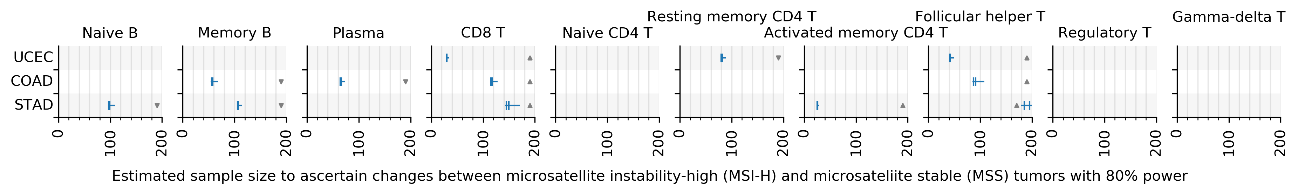

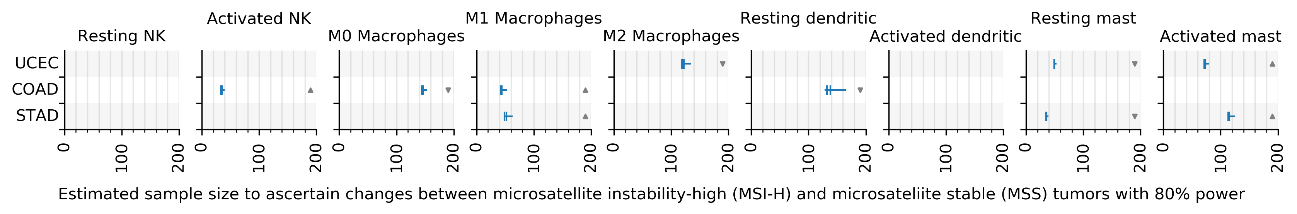

f

e

d

c

b

a

**Supplementary Figure 7 related to Fig. 3.** Sample size estimation by Sensei. **a,b** Estimated sample size for primary tumor vs recurrent tumor. More cell types are shown to supplement Fig. 3a. Estimations for unpaired test and paired test are shown in blue and yellow, respectively. Estimations are for infinite (left end of a whisker), 1,000 (left bar), 384 (right bar), and 100 (right end of a whisker) cells. Fewer cells require more samples to ascertain an effect. **c,d** Sample size estimation for normal tissue and tumor. More cell types are shown to supplement Fig. 3b. **e,f** Sample size estimation for MSI-H and MSS tumor. More cell types are shown to supplement Fig. 3c. Sample sizes over 200 are omitted.

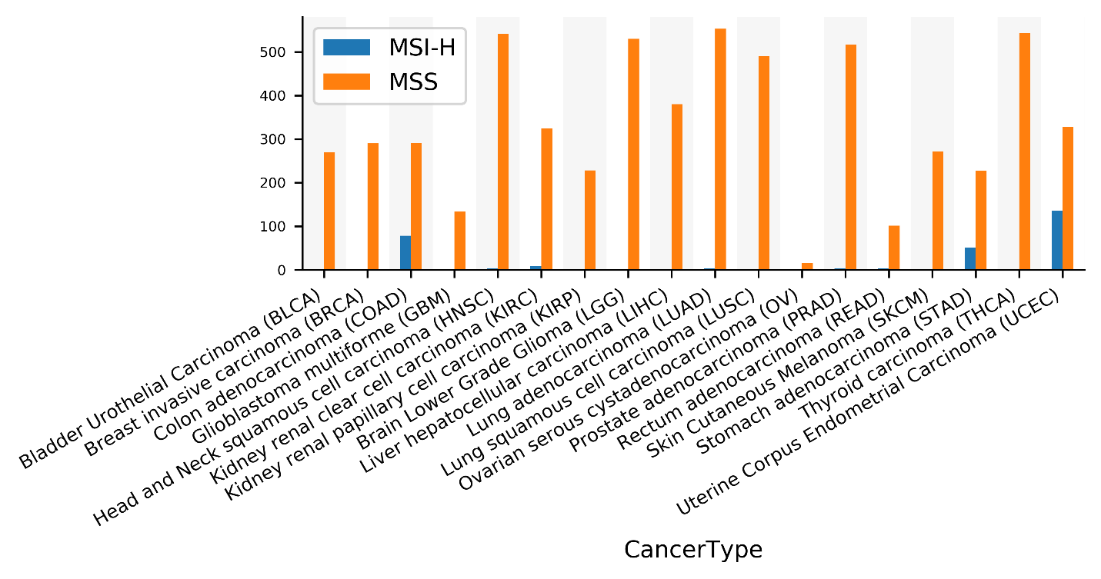

a

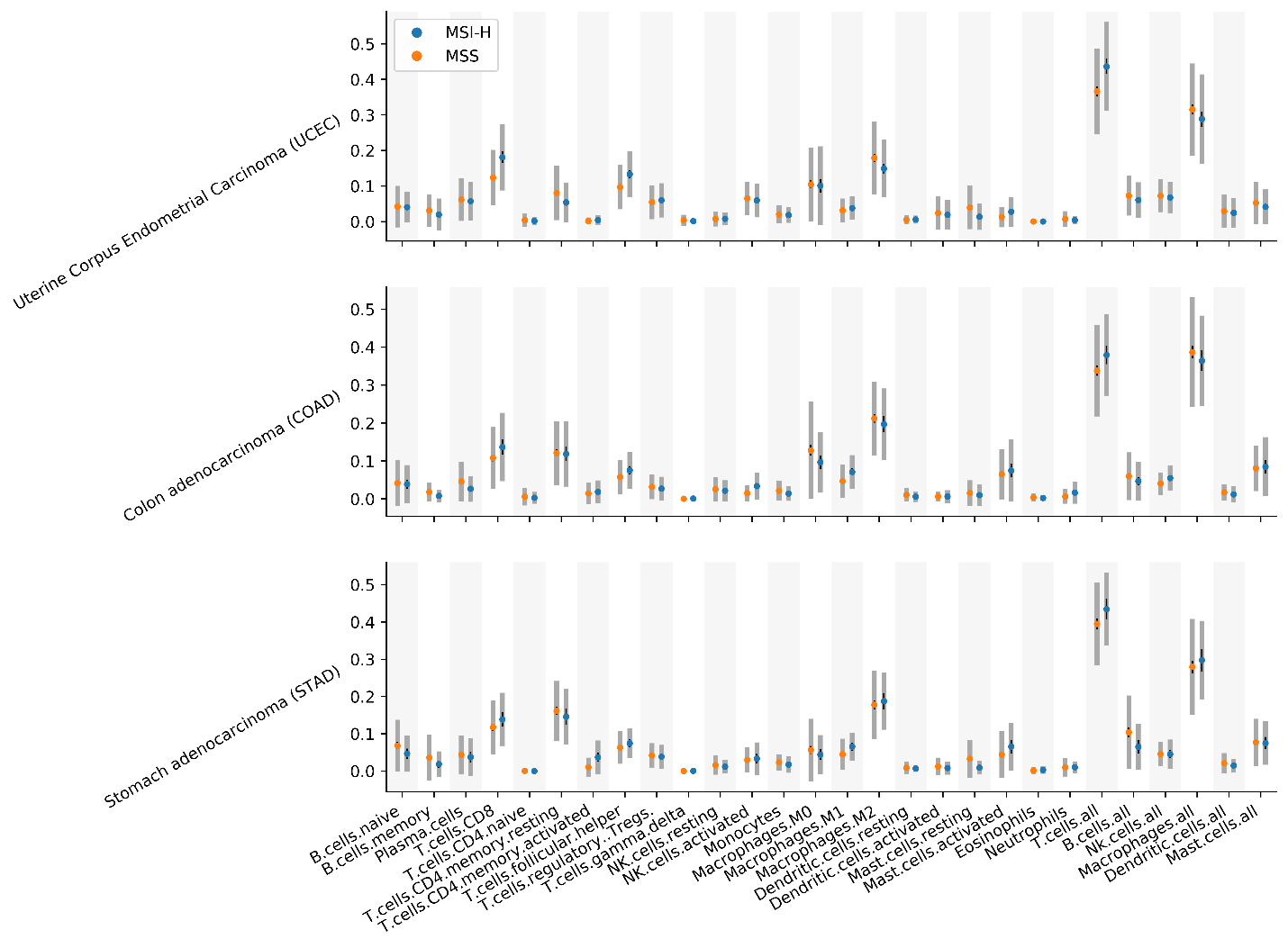

b

**Supplementary Figure 8. a** Number of MSI-H and MSS samples in each cancer type. **b** Immune cell type abundance in MSI-H and MSS tumor samples in UCEC, COAD, and STAD. Sample mean (dot), sample standard deviation (thick gray line spans a total of two times the standard deviation), and 95% confidence interval (think black line) of sample mean are shown.

| 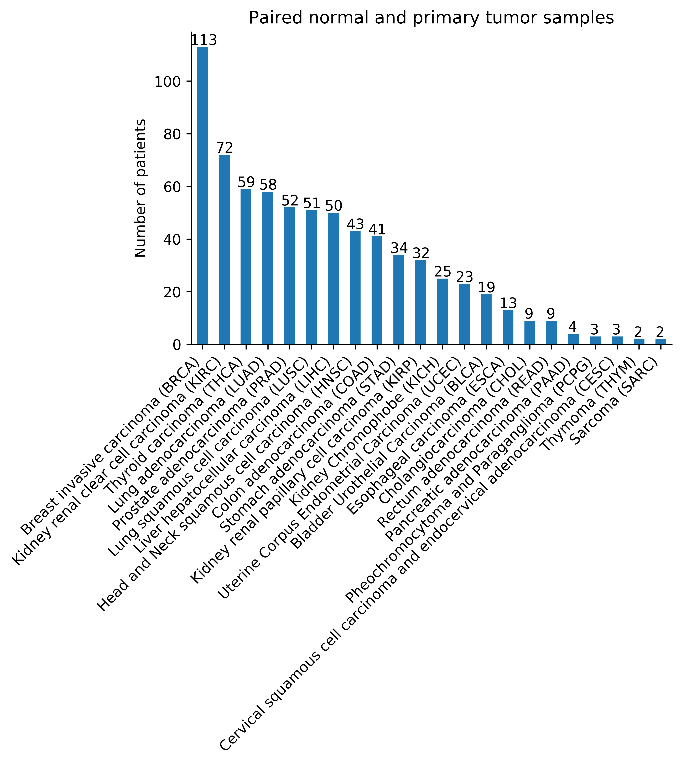  a | 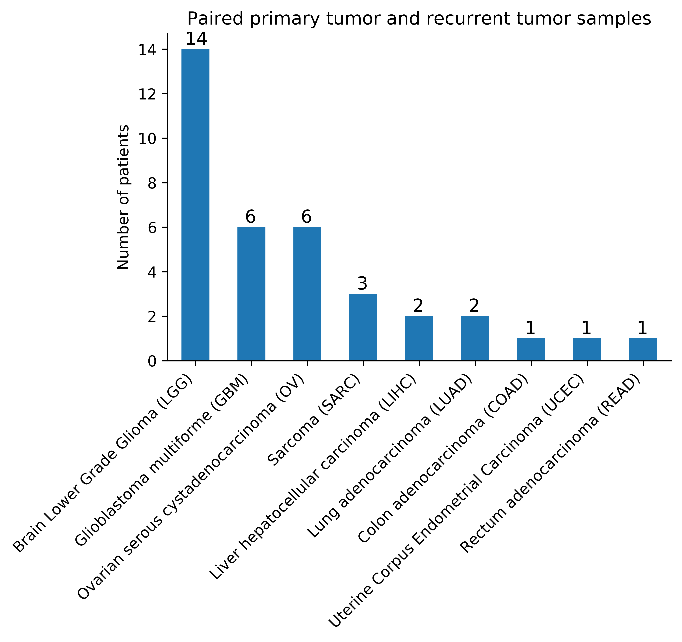  b |
| --- | --- |
| 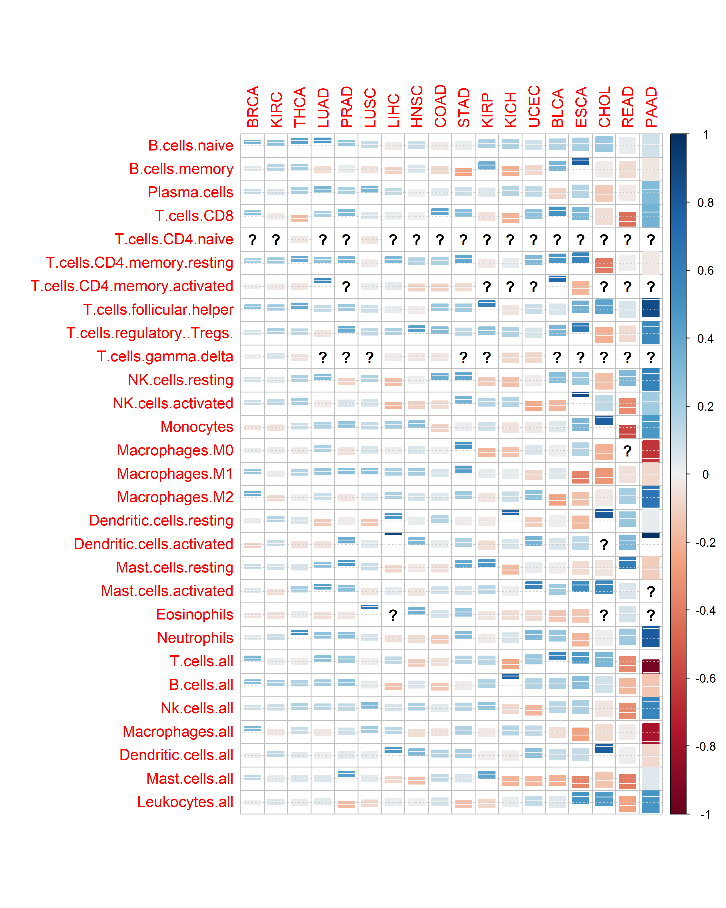  c | 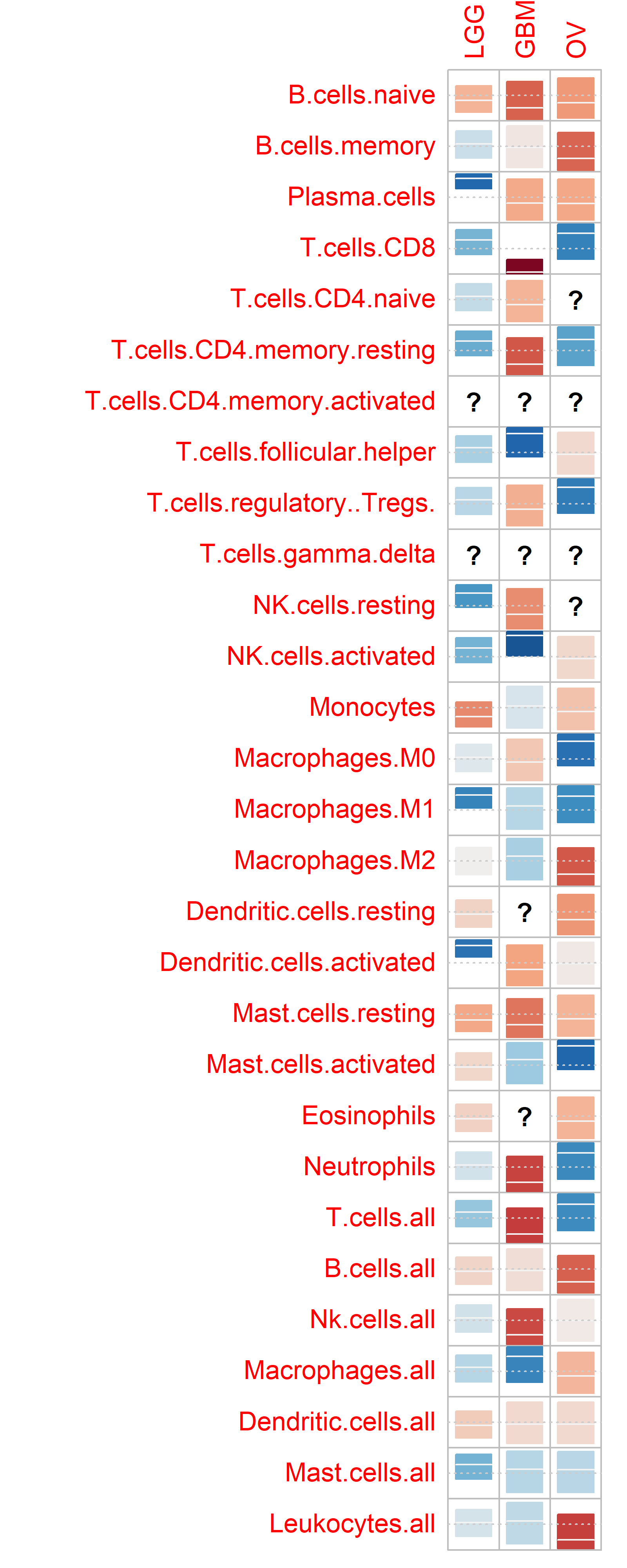  d |
| **Supplementary Figure 9.** Correlation of cell type abundances in paired samples in cancer data. **a** Number of patients with paired normal and primary tumor samples in each cancer type in the deconvolved TCGA dataset. **b** Number of patients with paired primary tumor and recurrent tumor samples in each cancer type. Cancer types with no paired samples do not show. **c** Correlation (center line/color: correlation; rectangle: 95% confidence interval; question marks: insufficient unique datapoint for calculating the correlation) of immune cell type abundances between paired normal and primary tumor samples in each cancer type. Cancer types with less than 4 patients are omitted. **d** Correlation of immune cell type abundances between paired primary tumor and recurrent tumor samples in each cancer type. | |

| 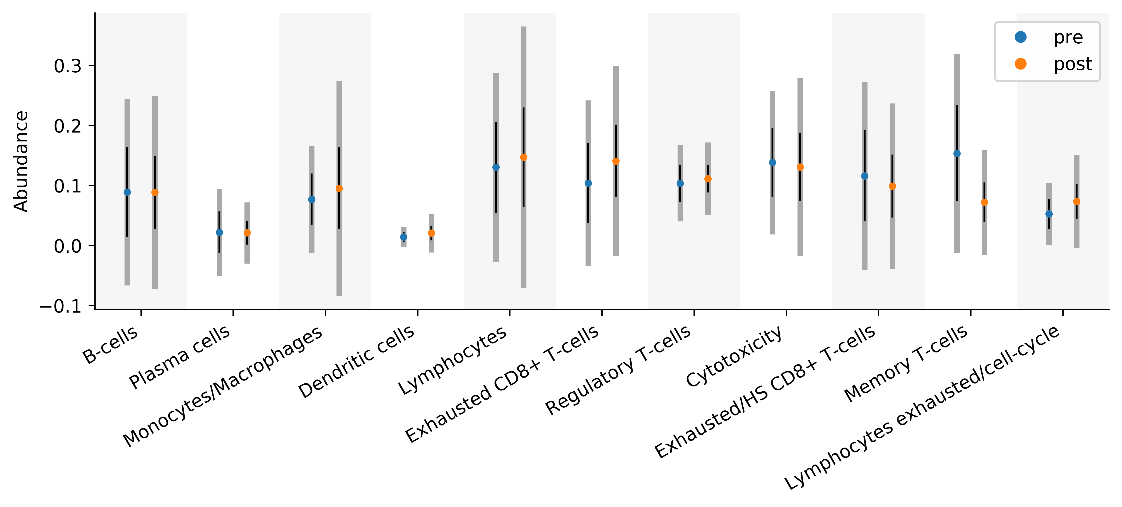  a | 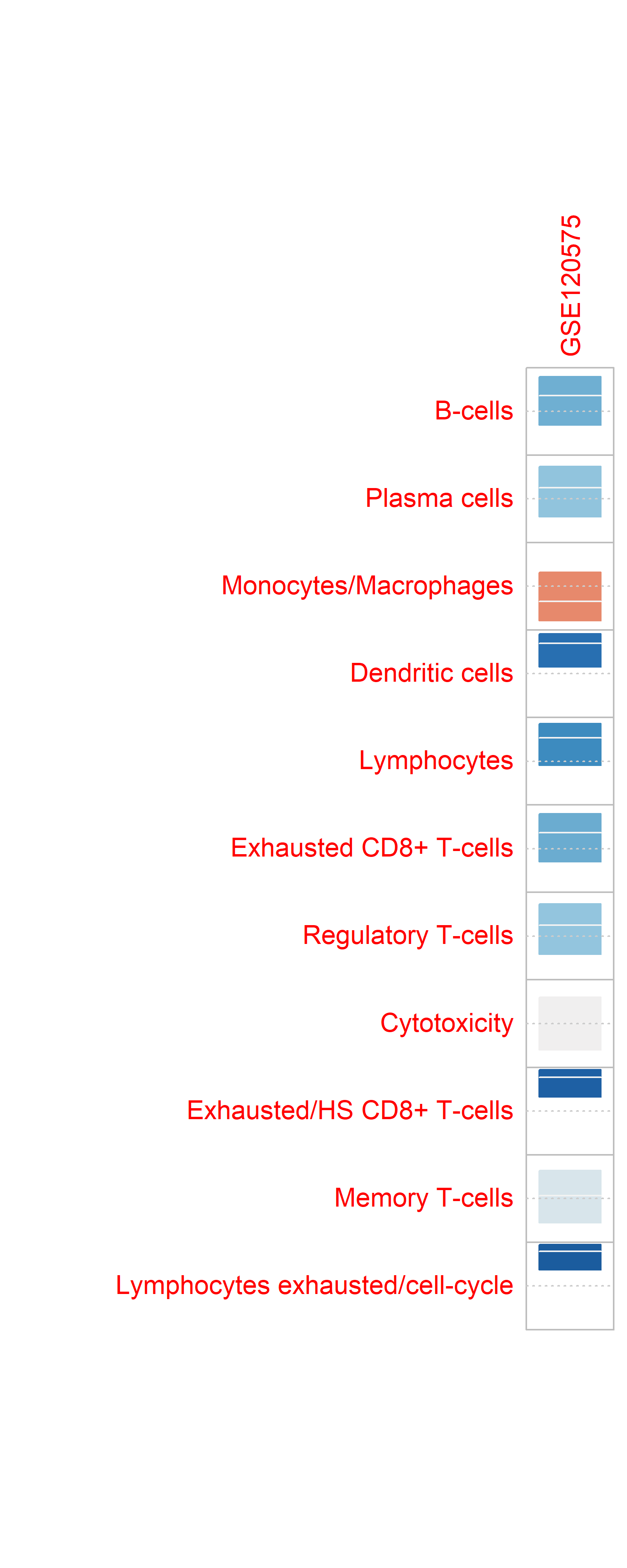  b |
| --- | --- |
| **Supplementary figure 10.** **a** Mean (dot), 95% confidence interval of mean (thin black line), and standard deviation (wide gray line) of the abundance of each major immune cell types in the metastatic melanoma data. **b** Correlation of immune cell types from pre- and post-treatment samples from metastatic melanoma patients. | |

| 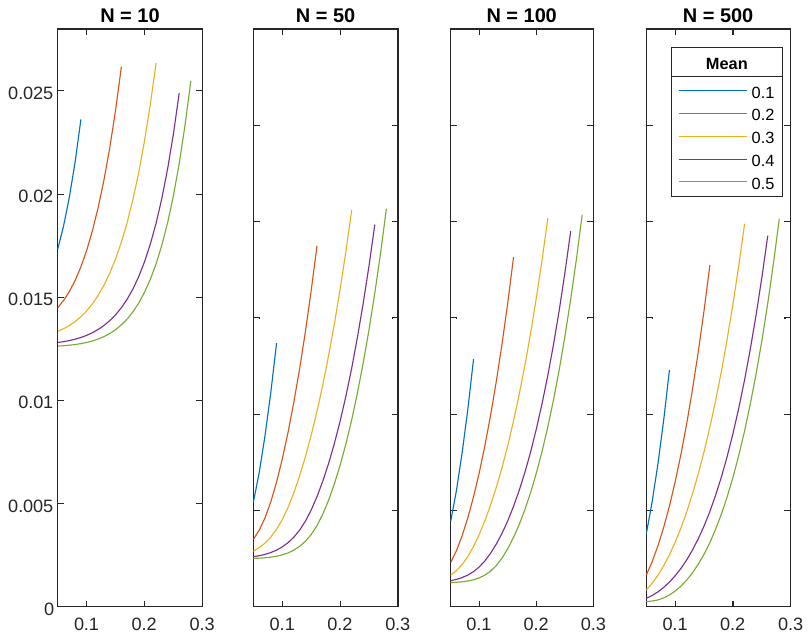  Error (L1 distance of CDF)  Standard deviation $\sigma$  $\mu$  a | 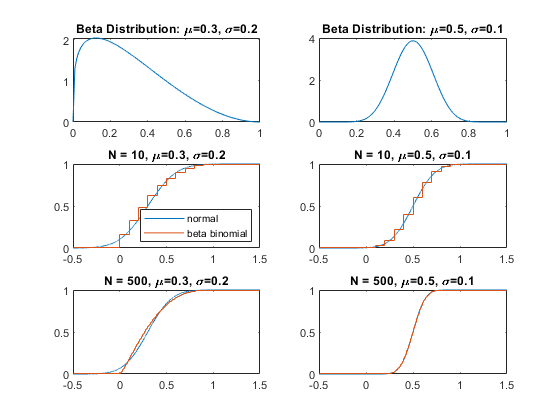  CDF  Density  CDF  b |
| --- | --- |
| **Supplementary Figure 11.** Comparison of the scaled beta-binomial distribution and the normal estimation. **a** L1 distance between the scaled beta-binomial distribution and normal distribution. Subpanel: number of cells. Color: mean; X-axis: standard deviation; Y-axis: L1 distance between cumulative distribution function (CDF) of a scaled beta binomial distribution and its normal approximation. **b** Four representative cases. First row: corresponding scaled beta-binomial distribution. Second row and third row: CDF for four cases. | |
