## Supplementary Text for "Sensei: How many samples to tell evolution in single-cell studies?"

### 1 Supplementary Text

#### 1.1 Validation of normal distribution

Skewness and kurtosis are standard measures for asymmetry and extreme values in a sample, respectively. Sample drawn from a normal distribution have skewness close to 0, and kurtosis also close to 0 (mesokurtic). Lower or higher skewness indicates left or right skew. Sample with lower or higher kurtosis are termed platykurtic or leptokurtic.

Because normality is an important assumption of the t-test, we verify it by calculating the skewness and kurtosis (Methods) of cell type abundances in TCGA data. Thorsson et al. deconvolved bulk RNA-seq data from TCGA data and obtained the proportion of immune cells [2]. Across 33 cancer types, available are 22 immune cell types, which we further grouped into 6 major types (T-cells, B-cells, NK cells, Macrophages, Dendritic cells, Mast cells). We calculated the skewness of each cell type in each sample, including normal tissues, primary tumors, and recurrent tumors. We found 90.5% of them between -3 and 3, and 78.0% between -2 and 2, indicating relatively good normality [28]. When considering the subtypes, as the abundances are closer to 0, distributions are naturally right-skewed. Nevertheless, 68.9% of the skewness is between -3 and 3, and 51.5% are between -2 and 2. The kurtosis, which is known to have less influence on the accuracy of the t-test [27], is also found to be within a reasonable range for most of the types. In specific, 87.5% and 66.3% of them are within -1 and 10 for major types and subtypes, respectively [28]. Overall, the t-test is proper for most of the immune cell types in most cancer types.

#### 1.2 User Manual for Sensei

We created a standalone web application using HTML and JavaScript. As Fig. 1 and Supplementary Figure 1 show, Sensei asks users to input a set of parameters for each group $i\in\{0,1\}$. Firstly, Sensei needs ranges of sample sizes $M_{i}$ for each group, which may factor in the financial feasibility and number of participants that can be gathered practically. Secondly, the number of cells in each single-cell sample, $N_{i}$, is needed to account for the variation introduced by limited/finite number of cells. Then, Sensei asks for the proportion of the cell type of interest in each group. Because cell type proportions vary among participants (in the same group) and may further be confounded by abovementioned technical variations, the proportion is not a deterministic value, but a random variable drawn from a beta distribution, which can be set by providing the mean $\mu$ and variance $\sigma$, or if preferred the standard parameters $\left( a_{i},b_{i} \right)$ for a beta distribution. These parameters may be obtained by surveying the literature, or by performing a pilot study, and a general guideline generated from TCGA is provided below. The graph at the top-right corner visualizes the beta distribution to double check if the distribution is satisfying. Finally, the user needs to set a false positive rate $\alpha$for t-test, usually 0.05 or 0.01, and a desired false negative rate $\beta$. For example, in Fig. 1g,h, Sensei generates a table of $\beta$ for $M_{0},M_{1}=8\sim12,$ $N_{0},N_{1}=1,000$, $\mu_{0}=0.3, \sigma_{0}=0.15, \mu_{1}=0.5, \sigma_{1}=0.1$, and false positive rate for t-test $\alpha=0.05$. A quick step-by-step guide is available at the webpage.

#### 1.3 Uncertainty in the estimated sample size

Uncertainty in the prior knowledge will propagate the estimated sample size. To reflect the uncertainty, a “confidence interval” can be constructed. When variances are fixed, the sample size is largely depending on the difference of the mean proportions of two groups. Thus, we calculate the confidence interval of the difference of the means. Using the Welch-Satterthwaite method, for two independent samples {$x_{0i}\}, i=1\ldots n_{0}$ and {$x_{1i}\}, i=1\ldots n_{1}$, the confidence interval $[\Delta\bar{x}_{L},\Delta\bar{x}_{U}]$ for $\Delta\bar{x}=\bar{x}_{1}-\bar{x}_{0}$ is constructed as

$$\begin{aligned} \Delta\bar{x}_{L},\Delta\bar{x}_{U}=\Delta\bar{x}\pm t_{1-\alpha/2,\nu}\sqrt{\frac{s_{0}^{2}}{n_{0}}+\frac{s_{2}^{2}}{n_{1}}},\#\left( 1 \right) \end{aligned}$$

where $(1-\alpha)$ is the confidence level and the degree of freedom

$$\begin{aligned} \nu=\frac{\left( \frac{s_{0}^{2}}{n_{0}}+\frac{s_{1}^{2}}{n_{1}} \right)^{2}}{\frac{s_{0}^{4}}{n_{0}^{2}(n_{0}-1)}+\frac{s_{1}^{4}}{n_{1}^{2}(n_{1}-1)}}.\#\left( 2 \right) \end{aligned}$$

We override the difference in the last step of sample size estimation. Note that if any of $\Delta\bar{x}$, $\Delta\bar{x}_{L}$, and $\Delta\bar{x}_{U}$ are of different signs, we consider the results unreliable.
